# LOCOM: A logistic regression model for testing differential abundance in compositional microbiome data with false discovery rate control

**DOI:** 10.1101/2021.10.03.462964

**Authors:** Yingtian Hu, Glen A. Satten, Yi-Juan Hu

## Abstract

**Motivation:** Compositional analysis is based on the premise that a relatively small proportion of taxa are “differentially abundant”, while the ratios of the relative abundances of the remaining taxa remain unchanged. Most existing methods of compositional analysis such as ANCOM or ANCOM-BC use log-transformed data, but log-transformation of data with pervasive zero counts is problematic, and these methods cannot always control the false discovery rate (FDR). Further, high-throughput microbiome data such as 16S amplicon or metagenomic sequencing are subject to experimental biases that are introduced in every step of the experimental workflow. McLaren, Willis and Callahan [1] have recently proposed a model for how these biases affect relative abundance data.

**Methods:** Motivated by [1], we show that the (log) odds ratios in a logistic regression comparing counts in two taxa are invariant to experimental biases. With this motivation, we propose LOCOM, a robust logistic regression approach to compositional analysis, that does not require pseudocounts. We use a Firth bias-corrected estimating function to account for sparse data. Inference is based on permutation to account for overdispersion and small sample sizes. Traits can be either binary or continuous, and adjustment for continuous and/or discrete confounding covariates is supported.

**Results:** Our simulations indicate that LOCOM always preserved FDR and had much improved sensitivity over existing methods. In contrast, ANCOM often had inflated FDR; ANCOM-BC largely controlled FDR but still had modest inflation occasionally; ALDEx2 generally had low sensitivity. LOCOM and ANCOM were robust to experimental biases in every situation, while ANCOM-BC and ALDEx2 had elevated FDR when biases at causal and non-causal taxa were differentially distributed. The flexibility of our method for a variety of microbiome studies is illustrated by the analysis of data from two microbiome studies.

**Availability and implementation:** Our R package LOCOM is available on GitHub at https://github.com/yijuanhu/LOCOM in formats appropriate for Macintosh or Windows.

## Background

Microbiome association studies are useful for the development of microbial biomarkers for prognosis and diagnosis of a disease or for the development of microbial targets (e.g., pathogenic or probiotic bacteria) for drug discovery, by detecting the taxa that are most strongly associated with the trait of interest (e.g., a clinical outcome or environmental factor). Read count data from 16S amplicon or metagenomic sequencing are typically summarized in a taxa count (or feature) table. Because the total sample read count (library size) is an experimental artifact, only the relative abundances of taxa, not absolute abundances, can be measured. Thus, microbial data are compositional (constrained to sum to 1). Analysis of microbial associations is further encumbered by data sparsity (having 50–90% zero counts in the taxa count table), high-dimensionality (having hundreds to thousands of taxa), and overdispersion. In addition, most microbiome association studies have relatively small sample sizes; further complications arise as the traits of interest may be either binary or continuous, and the detected associations may need to be adjusted for confounding covariates. Finally, any method for detecting taxon-trait associations should control the false discovery rate (FDR) [4]. The capability to handle all these features is essential for any statistical method to be practically useful.

There are (at least) two biological models for how microbial communities may change when comparing groups with different phenotypes or along a phenotypic gradient. In one model, a substantial proportion of the taxa in the community change; the concept “community state types” exemplifies this approach (see e.g., [5, 6]). The null hypothesis of “no differential abundance” that is tested at a taxon is that the taxon relative abundance remains the same, i.e., any change in taxon relative abundance across conditions is of interest. Methods for testing this hypothesis include metagenomeSeq [7] and the LDM [8]. In the other model, only a few key taxa are considered to change, while the other taxa show changes in relative abundance because of the compositional constraint. Thus, the null hypothesis is that the *ratios* of the relative abundances of the other taxa are unchanged. Methods for testing this hypothesis include ANCOM [9], ANCOM-BC [10], ALDEx2 [11], WRENCH [12], and DACOMP [13]. Because the hypothesis in the second model accounts for the compositional constraint that a change in relative abundance for one taxon necessarily implies a counterbalancing change in other taxa, it is generally referred to as *compositional analysis* [14].

Methods for compositional analysis are typically based on some form of log-ratio transformation of the read count data. The ratio can be formed against a reference taxon or the geometric mean of relative abundances of all taxa, referred to as additive log-ratio (alr) or centered log-ratio (clr) transformation, respectively [15]. Thus, zero count data, which cannot be log-transformed, is the major challenge in using compositional methods on microbiome data. A common practice is to add a *pseudocount*, most frequently 1 or 0.5 or even smaller values, to the zeros or all entries of the taxa count table [7, 9, 10, 15–17]. However, there is no consensus on how to choose the pseudocount, and it has been shown that the choice of pseudocount can affect the conclusions of a compositional analysis [18, 19].

The most popular pseudocount-based method for compositional analysis is perhaps ANCOM [9], which has now evolved into ANCOM-BC [10]. After adding 0.001 to all count data, ANCOM performs the alr transformation and treats the transformed data as the response of the linear regression model that includes the traits of interest and confounding variables as covariates. For each taxon, ANCOM uses all other taxa, one at a time, as the reference in forming the alr transformation, and then it employs a heuristic strategy to declare taxa that are significantly differentially abundant (outputting rankings of taxa instead of *p*-values). ANCOM-BC first estimates sampling fractions that are different across samples, and then models the log of read count data, in which zeros are replaced by pseudocount 1, through a linear regression model including the estimated sampling fraction as an offset term. This is essentially a normalization approach that first attempts to recover the absolute abundances of taxa and then test hypotheses about the absolute abundances. Unlike ANCOM, ANCOM-BC provides *p*-values for individual taxa. Both ANCOM and ANCOM-BC are restricted to group comparisons and can not handle continuous traits of interest, although adjustment for confounding covariates is supported.

Several methods have been developed that circumvent the use of pseudocount. ALDEx2 [11] first draws Monte-Carlo samples of non-zero relative abundances from Dirichlet distributions (with parameters constructed from read count data plus a uniform prior 0.5). Then, the sampled relative abundances are clr transformed and tested against the traits of interest via linear regression to yield *p*-values and adjusted *p*-values by the Benjamini-Hochberg (BH) procedure [20], both of which are averaged over sampling replicates to give the final *p*-values and adjusted *p*-values. In our simulations, we found that ALDEx2 tends to have low power, possibly due to the noise introduced in the sampling process. DACOMP [13] is a normalization approach that first selects a set of null reference taxa by a data-adaptive procedure and then normalizes read count data by rarefaction so that each taxon within the reference has similar counts across samples. However, the selected reference set may mistakenly contain causal taxa, which may compromise the performance of the normalization. In addition, adjustment for confounding covariates is not supported, although continuous traits of interest are allowed. WRENCH [12] is also a normalization approach that estimates group-specific compositional factors to bring the read counts of null taxa across groups to a similar level and employs DESeq2 to detect differentially abundant taxa. It is limited to group comparisons without confounding covariates.

It is also of interest to test differential abundance at the community (i.e., global) level, rather than taxon by taxon, using the compositional analysis approach. The most commonly used method for testing community-level hypotheses about the microbiome is PERMANOVA [21], which is a distance-based version of ANOVA. In the context of compositional analysis, the Aitchison distance can be used [14], which is simply the Euclidean distance applied to the clr transformed data [22]. Again, the clr transformation necessitates the use of pseudocount, so the choice of pseudocount may affect the outcome of the test.

Finally, it is of particular interest to develop a method that can provide valid inference even in the presence of experimental bias. Experimental bias is ubiquitous because each step in the sequencing experimental workflow (i.e., DNA extraction, PCR amplification, amplicon or metagenomic sequencing, and bioinformatics processing) preferentially measures (i.e., extracts, amplifies, sequences, and bioinformatically identifies) some taxa over others [1, 23–25]. For example, bacterial species differ in how easily they are lysed and therefore how much DNA they yield during DNA extraction [26]. As a result, the bias distorts the *measured* taxon relative abundances from their *actual* values.

We are particularly interested in the case of differential bias, where the bias of taxa that are associated with a trait is systematically different from the bias of null taxa. A concrete example of this is the differential bias between bacteria in the phyla *Bacteroidetes* and *Firmicutes. Bacteroidetes* are gram-negative, while *Firmicutes* are gram-positive. It is known that gram-positive bacteria have strong cell walls and are hence harder to lyse than gram-negative bacteria; thus gram-positive bacteria may be underrepresented due to bias in the extraction step of sample processing. The *Bacteroidetes*-*Firmicutes* ratio has been implicated in a number of studies of the gut microbiome (e.g., [27, 28]). Thus, studies that compare *Bacteroidetes* to *Firmicutes* may be affected by differential extraction bias. In some of our simulations, we consider the effect this kind of differential bias can have on the FDR.

In this article, we develop a novel method for compositional analysis of differential abundance, at both the taxon level and the global level, based on a robust version of logistic regression that we call LOCOM (LOgistic COMpositional). Our method circumvents the use of pseudocount, does not require the reference taxon to be null, and does not require normalization of the data. Further, it is applicable to a variety of microbiome studies with binary or continuous traits of interest and can account for potentially confounding covariates. In the methods section, we give the motivation for using logistic regression as a way to minimize the effect of experimental bias in analyzing microbiome data, and describe the details of our approach. In the results section, we present simulation studies that compare the performance of LOCOM to other compositional methods. We also compare results from LOCOM and other methods in the analysis of two microbiome datasets. We conclude with a discussion section.

## Methods

Let *Y*_*ij*_ be the read count of the *j*th taxon (*j* = 1, …, *J*) in the *i*th sample (*i* = 1, …, *n*) and *N*_*i*_ the library size of the *i*th sample. We denote by *P*_*ij*_ the observed relative abundance, given by *Y*_*ij*_*/N*_*i*_. We let *X*_*i*_ be a vector of *q* covariates including the (possibly multiple) traits of interest and other (confounding) covariates that we wish to adjust for, but excluding the intercept.

### Motivation

Our starting point is the model of McClaren, Willis and Callahan [1], as expanded by Zhao and Satten [29], which relates the expected value of the observed relative abundance, denoted by *p*_*ij*_, to the true relative abundance we would measure in an experiment with no experimental bias, denoted by *π*_*ij*_. In particular, this model assumes that

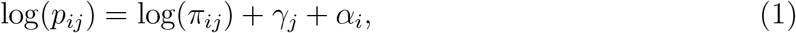

where *γ*_*j*_ is the taxon-specific *bias factor* that describes how the relative abundance is distorted by the bias, and *α*_*i*_ is the sample-specific *normalization factor* that ensures the composition constraint 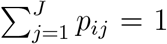. Following [29], we further assume that the true relative abundance *π*_*ij*_ can be described by a baseline relative abundance 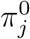 that would characterize the true relative abundance of taxon *j* for a sample having *X*_*i*_ = 0 and a term that describes how the baseline relative abundance is changed in the presence of covariates *X*_*i*_ ≠ 0. Then, we can replace (1) by

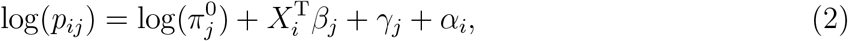

where *β*_*j*_ describes the way the true relative abundance changes with covariates *X*_*i*_ and is our parameter of interest. The presence of bias factors in (1) and (2) imply that inference based on the observed relative abundances *P*_*ij*_ may not give valid inference on *β*_*j*_. It is clear that, without knowing the bias factor *γ*_*j*_, we cannot estimate 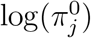 as 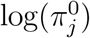 and *γ*_*j*_ always appear together as a sum.

We can examine equation (2) to see if there are any combinations of parameters that could potentially be estimated without knowing the bias factors. Analyzing log (probability) ratios such as log(*p*_*ij*_*/p*_*ij*′_) removes the effect of *α*_*i*_ (which depends on bias factors through normalization) but does not remove the effect of *γ*_*j*_. However, if we use (2) to write odds ratios of observed relative abundances for two different taxa and two different samples, we find

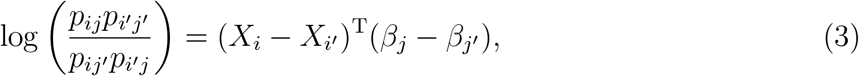

which *is* independent of bias factors. This motivates the choice of logistic regression to analyze microbiome count data.

Note that testing *β*_*j*_ − *β*_*j*′_ = 0 in (3) corresponds to testing *p*_*ij*_*/p*_*ij*′_ = *p*_*i*′*j*_*/p*_*i*′*j*′_, which is exactly the null hypothesis in a compositional analysis, e.g., in popular compositional models of the microbiome such as ANCOM and ALDEx2. As a result, logistic regression based on (3) is of interest even without the bias-removal motivation provided here.

### Multivariate logistic regression model

Equation (3) implies a polychotomous logistic regression of the full *n* × *J* taxa count table. This is numerically difficult as the analysis of each taxon potentially requires all *β*_*j*_ parameters. Instead, we follow Begg and Grey [30] and analyze data using individualized logistic regression of just two taxa at a time. Rather than considering all possible pairs of taxa, we choose one taxon (without loss of generality, the *J*th taxon) to be a reference taxon, and compare all other taxa to the reference taxon. Then, if we define *µ*_*ij*_ = *p*_*ij*_/(*p*_*ij*_ + *p*_*iJ*_), equation (2) implies

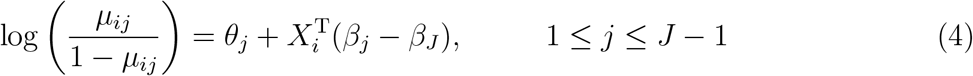

where the intercept 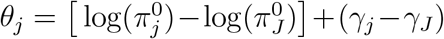 is treated as a free, nuisance parameter. The model is over-parameterized and only *J* − 1 of (*β*_1_, *β*_2_, …, *β*_*J*_) are identifiable. We set *β*_*J*_ = 0 to ensure identifiability. According to [30], the efficiency of individualized logistic regression highly depends on the prevalence (relative abundance) of the reference category, so we recommend that the reference taxon be a common taxon that is present in a large number of samples.

To avoid distributional assumptions in a standard logistic regression, we consider the score functions as estimating functions. When a taxon is rare and/or the sample size is small, it may occur that all (or nearly all) counts for that taxon are zero in one group (e.g., the case or control group), which is referred to as separation in the literature on logistic regression. It is known that the Firth bias correction [2], when applied to logistic regression [3], solves the problem of separation. Hence, we estimate *β*_*j*_ by solving the Firth-corrected score equations

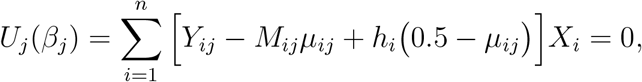

where *M*_*ij*_ = *Y*_*ij*_+*Y*_*iJ*_ and *h*_*i*_ is the *i*th diagonal element of the weighted hat matrix 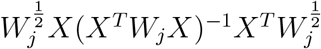 with the weight matrix *W*_*j*_ = Diag[*M*_*ij*_*µ*_*ij*_(1 − *µ*_*ij*_)]. We let 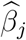 denote the estimator of *β*_*j*_ obtained by solving the above equation.

### Testing hypotheses at individual taxa

Now we derive the formula for the null hypotheses that correspond to null taxa. Write *β*_*j*_ = (*β*_*j*,1_, *β*_*j*,−1_), where *β*_*j*,1_ is the coefficient for the trait of interest and *β*_*j*,−1_ for the other covariates. We assume one trait of interest here although our methodology is readily generalizable to test multiple traits simultaneously. The naive formula *β*_*j*,1_ = 0 corresponds to a null taxon only when the reference taxon used in (4) is null. As we have no such knowledge about the reference taxon *a priori*, we seek a formula that does not require such knowledge; in addition, we need a test for the reference taxon *per se*.

To this end, we make the assumption that more than half of the taxa are null taxa, which has been frequently adopted in compositional methods [12, 13]. We use the formula

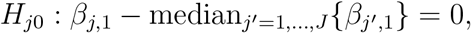

where *j* = 1, …, *J*. Recall that *β*_*J*,1_ = 0, which is included in the median calculation and also used to obtain a test for the reference taxon. With the assumption, we expect median_*j*′ =1,…,*J*_{*β*_*j*′, 1_} to correspond to the value of *β*_*j*′, 1_ for some null taxon *j*′. Thus, *H*_*j*0_ always corresponds to a test of taxon *j* against a null taxon, irrespective of whether the reference taxon *J* is null or not. Note that the clr transformation 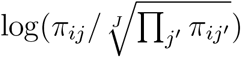 is equivalent to subtracting mean_*j*′ =1,…,*J*_{*β*_*j*′, 1_} off *β*_*j*,1_, but the mean is sensitive to large or outlying observations.

For testing *H*_*j*0_, it is natural to use the test statistic 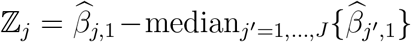. In the simplest case testing a binary trait with no other covariates, ℤ_*j*_ is invariant to different choices of the reference taxon, since all pairwise log odds ratios (*β*_*j*_ − *β*_*j*′_) in this case are estimated by the empirical log odds ratios log{*n*_1*j*_*n*_0*j*′_ /(*n*_0*j*_*n*_1*j*′_)}, where 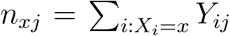. This holds even if the Firth-corrected estimator is used because, in this simple case, the Firth-corrected estimator corresponds to adding 1/2 to each *n*_*xj*_ [2, 3]. For general cases, we evaluate the dependence of ℤ_*j*_ on the reference taxon via simulations.

To avoid distributional assumptions in sparse microbiome data, we assess the significance of ℤ_*j*_ using the permutation scheme for logistic regression proposed by Potter [31]. Specifically, the covariate vector *X*_*i*_ is partitioned into (*T*_*i*_, *C*_*i*_) where *T*_*i*_ denotes the trait of interest and *C*_*i*_ the other covariates including the intercept. A linear regression of *T*_*i*_ on *C*_*i*_ is fit to obtain the residual *T*_*ir*_, which is then permuted to obtain 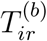 and to construct the new covariate vector 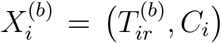. We follow the same procedure as for the observed dataset to obtain the estimate of *β*_*j*,1_ from the *b*th permutation replicate, denoted by 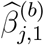, and the corresponding statistic 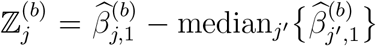. We adopt Sandve’s sequential stopping rule [32] with a minor modification to stop the permutation procedure, which is described below. At the *B*th *current* permutation, we record the numbers of times that 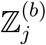 falls on the left and right hand side of ℤ_*j*_ by *L*_*j*_ and *R*_*j*_, respectively, and count the number of *rejection* to be 2 min(*L*_*j*_ + 1, *R*_*j*_ + 1). The current *p*-value is given by *p*_*j*_ = 2 min(*L*_*j*_ + 1, *R*_*j*_ + 1)/(*B* + 1) and the current *q*-value is calculated according to [32]. The permutation procedure is continued until each taxon either has the *q*-value below the nominal FDR level or has the number of rejection exceeding a pre-specified level (e.g., 100). This stopping rule is slightly different from Sandve’s in that we obtain 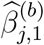 for every taxon at every permutation, rather than stopping permutation early for some taxa, because the median calculation requires 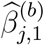 from all taxa.

### Testing the global hypothesis

The global null hypothesis is that there are no differentially abundant taxa, i.e., *H*_*j*0_ holds for every taxon. Given the *p*-values at individual taxa, it is straightforward to construct a global test statistic by combining the individual *p*-values. Here we adopt the harmonic-mean approach proposed by Wilson et al. [33] to combining *p*-values, which is more robust to the dependence structure among taxa than Fisher’s method. The harmonic mean of *p*_*j*_s is 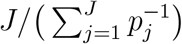, whose smaller values correspond to stronger evidence against the null hypothesis. To have a usual test statistic with a reversed directionality, we choose 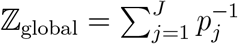. We use all permutation replicates generated for taxon-level tests, say *B*, to assess the significance of ℤ_global_. At the *b*th permutation replicate, the test statistic is 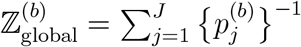, where 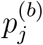 is the *p*-value of taxon *j* at this null replicate. Following [34], we calculate the null *p*-value 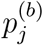 using the rank statistic to be 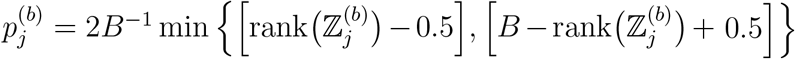, where rank 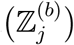 is the rank of 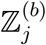 among *B* such statistics. Let *R*_global_ be the number of times that 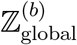 falls on the right hand side of ℤ_global_. Then, the global *p*-value is given by (*R*_global_ + 1) /(*B* + 1).

## Results

### Simulation studies

We used simulation studies to evaluate the performance of LOCOM and compare its performance to other currently-available compositional analysis packages. We based our simulations on data on 856 taxa of the upper-respiratory-tract (URT) microbiome; these taxa correspond to the “OTUs” in the original report on these data by Charlson et al. [35]. We considered both binary and continuous traits of interest and both binary and continuous confounders, as well as the case of no confounder. We assumed two causal mechanisms. For the first mechanism (referred to as M1), we randomly sampled 20 taxa (after excluding the most abundant taxon) whose mean relative abundances were greater than 0.005 as observed in the URT data to be *causal* (i.e., associated with the trait of interest). For the second mechanism (referred to as M2), we selected the top five most abundant taxa (having mean relative abundance 0.105, 0.062, 0.054, 0.050, and 0.049) to be *causal*. For simulations with a confounding covariate, we assumed the confounder was associated with 20 taxa under M1 (10 sampled at random from the 20 causal taxa and 10 from the null taxa) and 5 taxa under M2 (2 from the 5 causal taxa and 3 from the null taxa). We simulated most data without adding experimental bias, but did conduct one set of simulations having differential experimental bias. We focused on data sets having 100 observations but also considered some data sets with 200 observations.

To be specific, we let *T*_*i*_ denote the trait and *C*_*i*_ the confounder for the *i*th sample. To generate a binary trait, we selected an equal number of samples with *T*_*i*_ = 1 and *T*_*i*_ = 0. When a binary confounder was present, we drew *C*_*i*_ from the Bernoulli distribution with probability 0.2 in samples with *T*_*i*_ = 0 and from the Bernoulli distribution with probability 0.8 in samples with *T*_*i*_ = 1. When a continuous confounder was present, we drew *C*_*i*_ from the uniform distribution *U*[−1, 1] in samples with *T*_*i*_ = 0 and *U*[0, 2] in samples with *T*_*i*_ = 1. To generate a continuous trait, we sampled it from *U*[−1, 1] when there was no confounder. When there was a binary confounder, we used the aforementioned data generated for a binary trait and a continuous confounder but exchanged the roles of trait and confounder. When there was a continuous confounder, we generated *T*_*i*_ from *U*[−1, 1] and a third variable *Z*_*i*_ from *U*[−1, 1] independently of *T*_*i*_, and then constructed the confounder 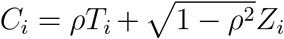, where *ρ* was fixed at 0.5.

To simulate read count data for the 856 taxa, we first sampled the *baseline* (when *T*_*i*_ = 0 and *C*_*i*_ = 0) relative abundances 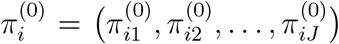 of all taxa for each sample from the Dirichlet distribution 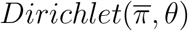, where the mean parameter 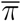 and overdispersion parameter *θ* took the estimated mean and overdispersion (0.02) in the Dirichlet-Multinomial (DM) model fitted to the URT data. We formed the relative abundances *p*_*ij*_ for all taxa by spiking the *j*′th causal taxon with an exp(*β*_*j*′, 1_)-fold change and the *j″*th confounder-associated taxon with an exp(*β*_*j″*, 2_)-fold change, then re-normalizing the relative abundances, so that

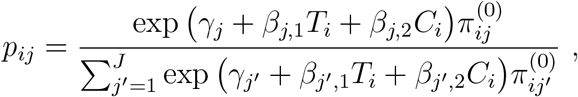

where *γ*_*j*_ was the bias factor for the *j*th taxon. Note that *β*_*j*,1_ = 0 for null taxa, *β*_*j*,2_ = 0 for confounder-independent taxa, and *γ*_*j*_ = 0 for all taxa for data without experimental bias. In most cases, for simplicity, we set *β*_*j*,1_ = *β* for all causal taxa, and thus *β* is a single parameter that we refer to as the effect size; we refer to exp(*β*) as the fold change. In some cases, we also considered the more general scenario when different values were sampled for different *β*_*j*,1_. We fixed *β*_*j*,2_ = log(2) for all confounder-associated taxa. When there was no confounder, we simply dropped the term *β*_*j*,2_*C*_*i*_ in calculating *p*_*ij*_. In cases with differential experimental bias, we drew *γ*_*j*_ from *N* (0, 0.8^2^) for non-causal taxa and from *N* (1, 0.8^2^) for causal taxa. Finally, we generated the taxon count data for each sample using the Multinomial model with mean *π*_*i*_ = (*π*_*i*1_, *π*_*i*2_, …, *π*_*iJ*_) and library size sampled from *N*(10000, (10000/3)^2^) and left-truncated at 2000.

We applied two versions of LOCOM: one used the most abundant null taxon as the reference, which is referred to as LOCOM-null, and one used the most abundant causal taxon as the reference, referred to as LOCOM-causal. In practice when the most abundant taxon is chosen as the reference, LOCOM-null would be used in M1 and LOCOM-causal in M2; the other version served as an internal check of the robustness of LOCOM to the choice of the reference taxon. For testing the global hypothesis, we compared LOCOM to PERMANOVA (the adonis2 function in the vegan R package) based on the Aitchison distance, which is referred to as PERMANOVA-half and PERMANOVA-one corresponding to adding pseudocount 0.5 and 1, respectively, to all cells. The type I error and power of the global test were assessed at the nominal level 0.05 based on 5000 and 1000 replicates of data, respectively. For testing individual taxa, we compared LOCOM to ANCOM, ANCOM-BC, ALDEx2, DACOMP, and WRENCH. However, ANCOM, ANCOM-BC, and WRENCH cannot handle continuous traits; DACOMP and WRENCH cannot adjust for other covariates. Prior to analysis, we removed taxa having fewer than 20% presence (i.e., present in fewer than 20% of samples) in each simulated dataset. For ANCOM and ANCOM-BC, we also considered their own filtering criterion with 10% presence as the cutoff and refer to these methods as ANCOM^*o*^ and ANCOM-BC^*o*^. In the case with a binary trait only, we considered two additional methods, Pseudo-half and Pseudo-one, which add pseudocount 0.5 and 1, respectively, to all cells, form the alr using the most abundant null taxon as the reference, perform the Wilcoxon rank-sum test at individual log ratios, and correct multiple comparisons using the Benjamini-Hochberg method. Because the reference was selected to be a taxon known to be null, these methods are not applicable to real studies but are included in the simulations here to assess the properties of the pseudocount approach to testing individual taxa. The sensitivity (proportion of truly causal taxa that were detected) and empirical FDR were assessed at nominal FDR 20% based on 1000 replicates of data. We chose a relatively high nominal FDR because the numbers of causal taxa in both M1 and M2 were small.

### Simulation results

The type I error of the global tests for all simulation scenarios are summarized in Table 1. In all scenarios, LOCOM-null and LOCOM-causal yielded type I error rates that were close to the nominal level and generally closer for sample size 200 than 100. Note that, in cases when there was a confounder, there was substantial inflation of type I error when the confounder was not accounted for (Table S1), demonstrating that LOCOM is effective in adjusting for confounders. The PERMANOVA tests also controlled type I error. In cases without any confounder, the zero data were similarly distributed across trait values under the (global) null, so the effect of adding pseudocount is non-differential. In cases with a confounder, the taxa associated with the confounder caused the zeros to be differentially distributed across trait values, so that adding pseudocount had a differential effect for different trait values; however, this difference was controlled by adding the confounder as a covariate in the model. Note that, although the pseudocount approach did not lead to invalid global tests, it did lead to invalid tests at individual taxa (in the presence of causal taxa), as indicated in the FDR of Pseudo-one and Pseudo-half (Figures 1 and Figures S3).

**Table 1:** Type I error for testing the global hypothesis at nominal level 0.05

| Trait | Confounder | Causal mechanism | $n$ | LOCOM<br>-null | LOCOM<br>-causal | PERMANOVA<br>-half | PERMANOVA<br>-one |
| --- | --- | --- | --- | --- | --- | --- | --- |
| Binary | NA | M1, M2 | 100 | 0.052 | 0.052 | 0.056 | 0.063 |
|  |  |  | 200 | 0.055 | 0.055 | 0.048 | 0.047 |
| Continuous | NA | M1, M2 | 100 | 0.049 | 0.049 | 0.059 | 0.055 |
|  |  |  | 200 | 0.049 | 0.053 | 0.050 | 0.050 |
| Binary | Binary | M1 | 100 | 0.060 | 0.037 | 0.049 | 0.049 |
|  |  |  | 200 | 0.050 | 0.045 | 0.050 | 0.050 |
|  |  | M2 | 100 | 0.040 | 0.059 | 0.053 | 0.052 |
|  |  |  | 200 | 0.043 | 0.052 | 0.048 | 0.049 |
| Binary | Continuous | M1 | 100 | 0.049 | 0.042 | 0.054 | 0.053 |
|  |  |  | 200 | 0.049 | 0.044 | 0.050 | 0.051 |
|  |  | M2 | 100 | 0.046 | 0.053 | 0.049 | 0.053 |
|  |  |  | 200 | 0.049 | 0.053 | 0.048 | 0.051 |
| Continuous | Binary | M1 | 100 | 0.048 | 0.048 | 0.045 | 0.045 |
|  |  |  | 200 | 0.049 | 0.051 | 0.051 | 0.050 |
|  |  | M2 | 100 | 0.047 | 0.049 | 0.047 | 0.045 |
|  |  |  | 200 | 0.053 | 0.052 | 0.049 | 0.047 |
| Continuous | Continuous | M1 | 100 | 0.055 | 0.049 | 0.048 | 0.048 |
|  |  |  | 200 | 0.049 | 0.051 | 0.050 | 0.051 |
|  |  | M2 | 100 | 0.043 | 0.047 | 0.048 | 0.048 |
|  |  |  | 200 | 0.053 | 0.052 | 0.050 | 0.051 |

Note: $n$ is the number of samples. When there was no confounder, the data generated under the global null of M1 and M2 were the same.

**Figure 1:**
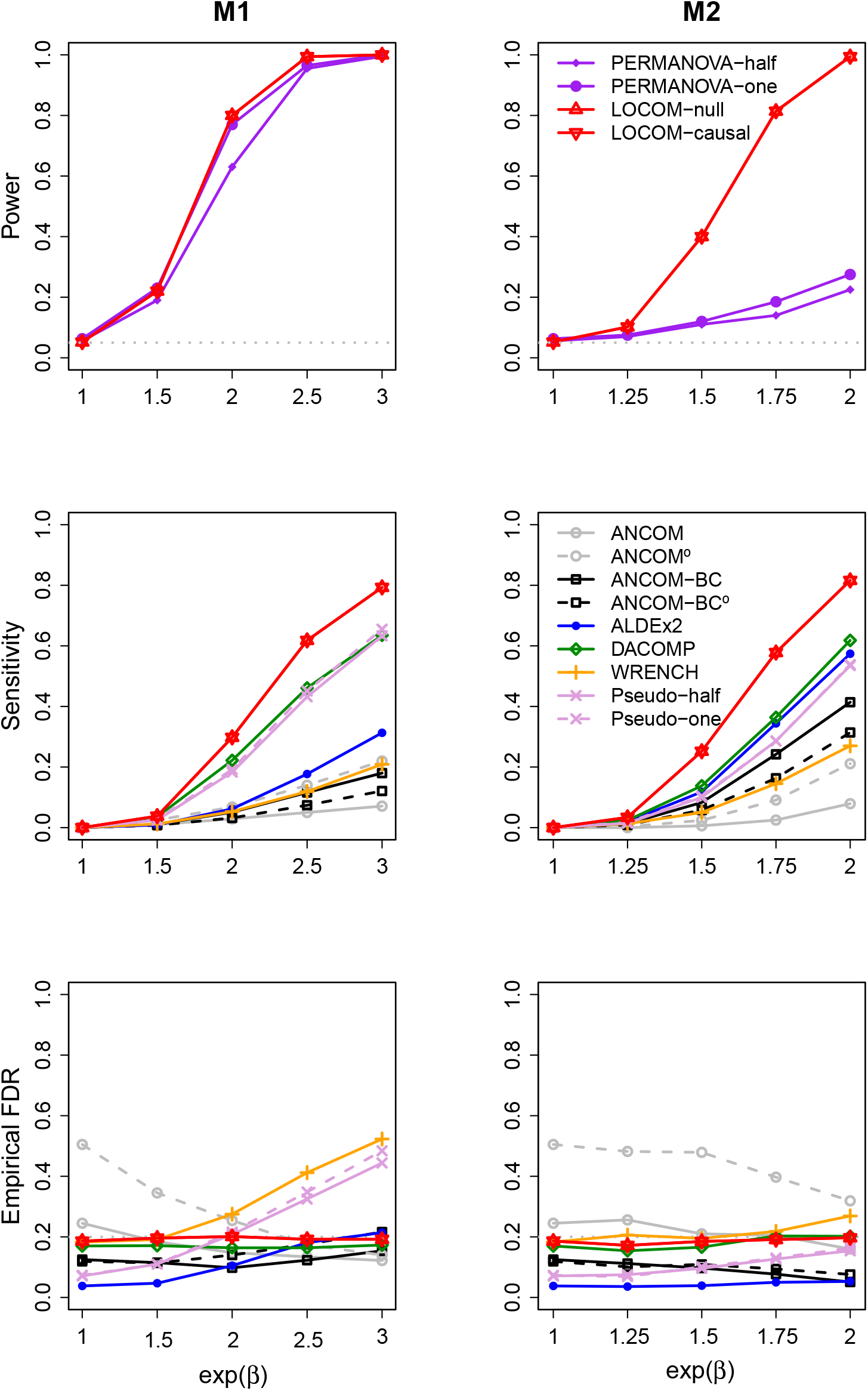
Simulation results for data (*n* = 100) with a binary trait (and no confounder). The power at exp(*β*) = 1 corresponds to the type I error. The gray dotted line indicates the nominal type I error 0.05 in the first row and the nominal FDR 20% in the last row.

Figures 1–4 present power of the global tests and sensitivity and empirical FDR of the individual taxon tests, for a binary or continuous trait without and with a binary confounder. The results for cases with a continuous confounder are deferred to Figures S1–S2, which show similar patterns of results to their counterparts with a binary confounder (Figures 3–4). While these figures pertain to the sample size 100, Figures S3–S8 pertain to the sample size 200 and show similar patterns of results to their counterparts with the sample size 100.

**Figure 2:**
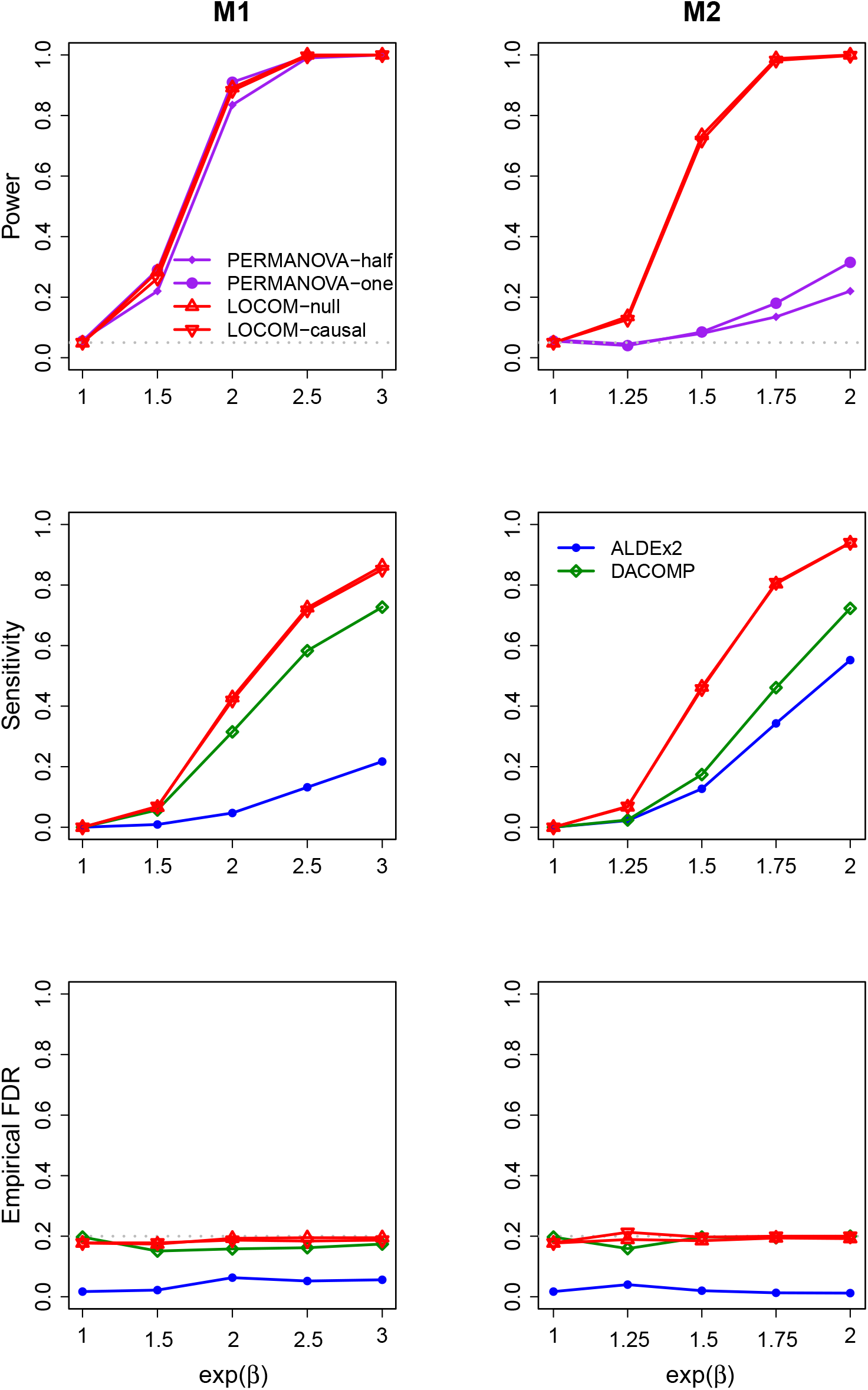
Simulation results for data (*n* = 100) with a continuous trait (and no confounder).

**Figure 3:**
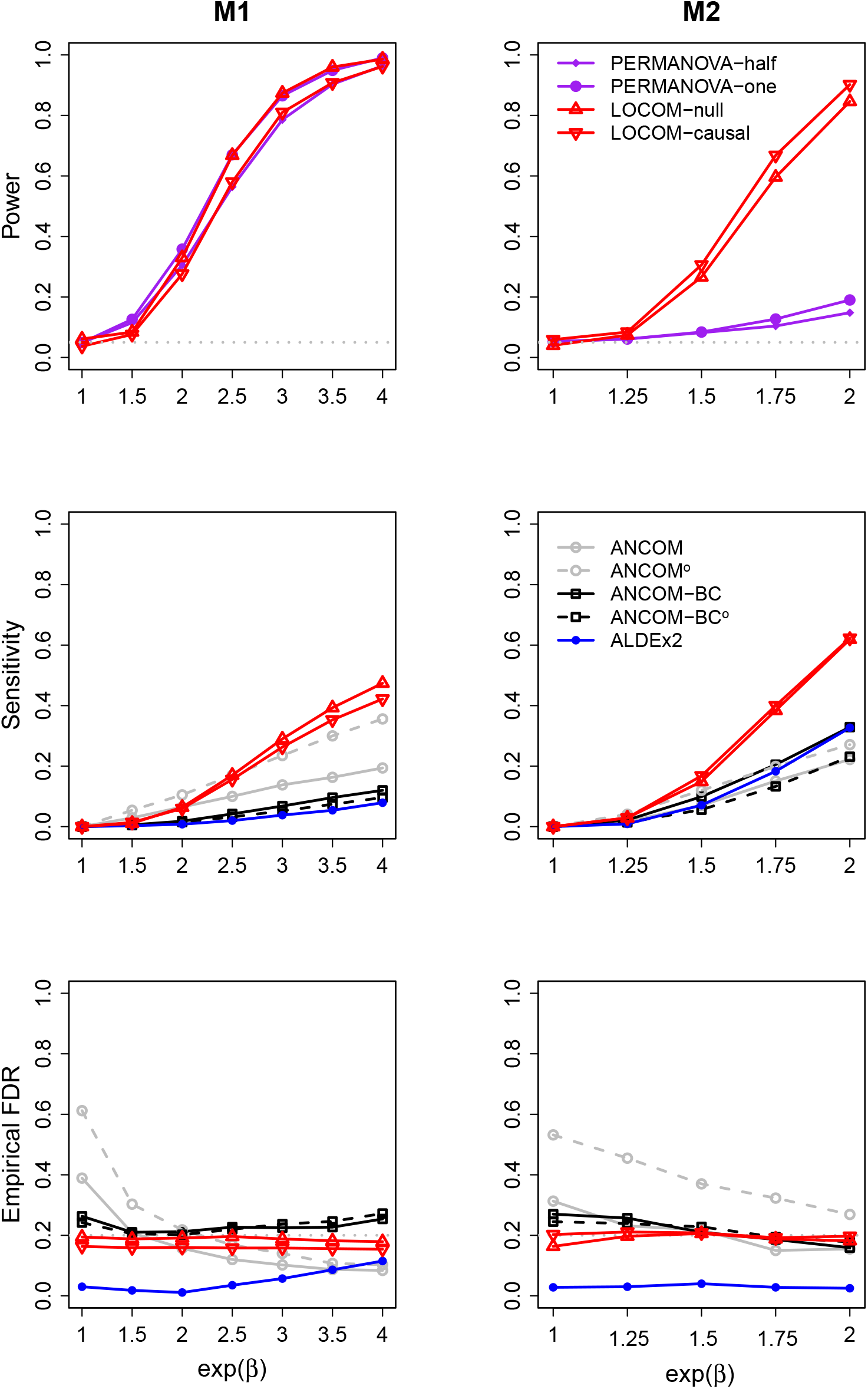
Simulation results for data (*n* = 100) with a binary trait and a binary confounder.

**Figure 4:**
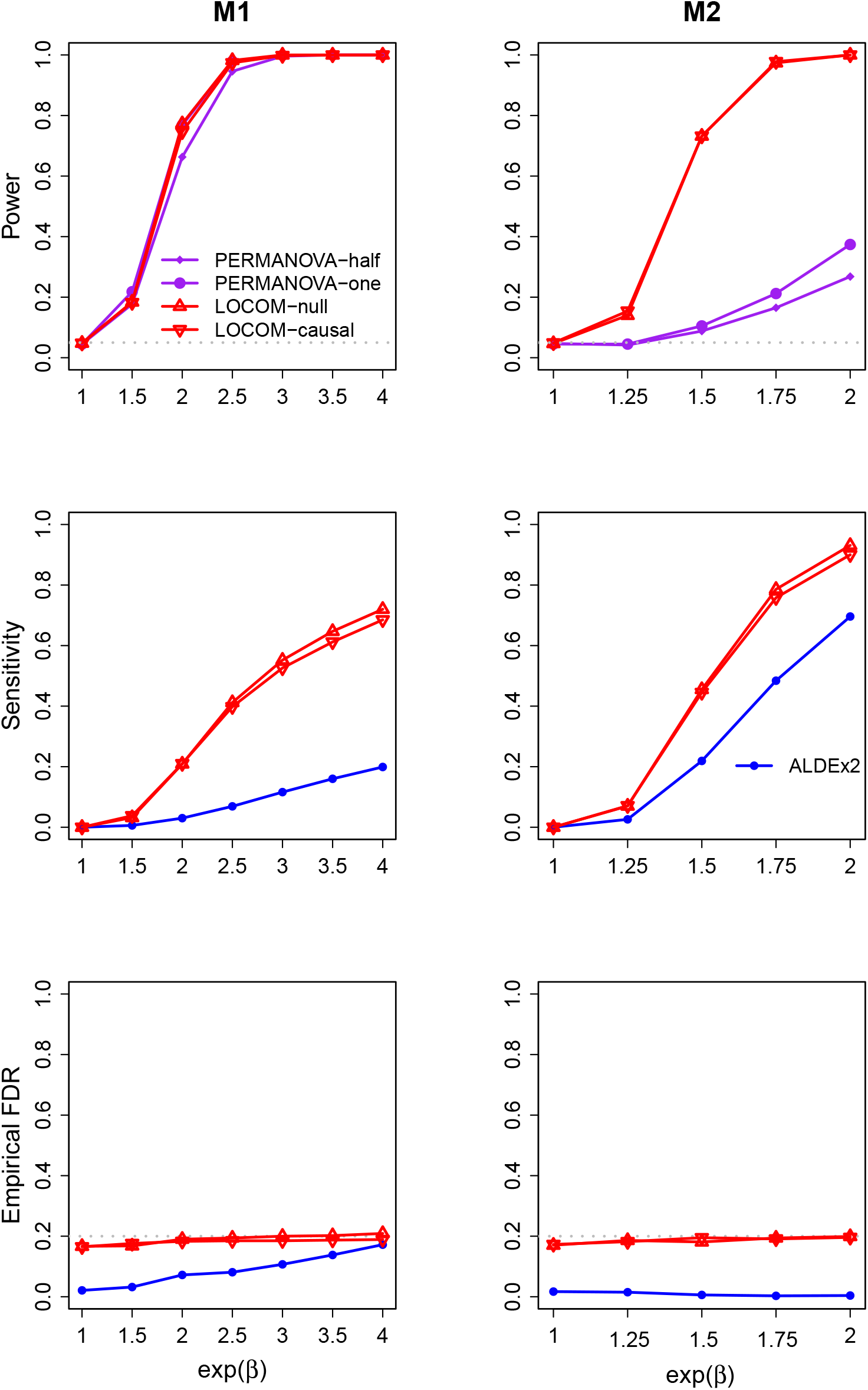
Simulation results for data (*n* = 100) with a continuous trait and a binary confounder.

In the simplest scenario with a binary trait and no confounder (Figures 1 and S3), LOCOM-null and LOCOM-causal yielded identical results; in fact, the two methods gave identical *p*-values for every dataset in this case, which corroborates our claim that the test is invariant to different reference taxa. In other scenarios, LOCOM-null and LOCOM-causal produced similar results although the one using the more abundant taxon as the reference (LOCOM-null in M1 and LOCOM-causal in M2) tended to be more powerful and more sensitive. In all scenarios, the LOCOM tests yielded the highest power for testing the global hypothesis and the highest sensitivity for testing individual taxa while always controlling the FDR.

The competing methods generally have limited application to the scenarios we considered and significantly inferior performance to LOCOM. PERMANOVA had similar power to LOCOM in M1 but lost substantial power to LOCOM in M2. For testing individual taxa, ALDEx2 is the only method that is applicable to all scenarios; although it controlled the FDR in most cases, it still lost control occasionally (S3 and S7) and it had much lower sensitivity than LOCOM in all cases. ANCOM and ANCOM-BC are only applicable for testing binary traits, with or without confounders. ANCOM easily lost control of FDR, especially with their own, less stringent filtering criterion. ANCOM-BC controlled the FDR better than ANCOM but still had some modest inflation (e.g., Figure 3). Both ANCOM and ANCOM-BC had substantially lower sensitivity than LOCOM when they controlled the FDR. DACOMP is applicable for testing both binary and continuous traits but without any confounder; in these scenarios, DACOMP largely controlled the FDR but still lost control occasionally (Figure S3, under M2); although the sensitivity of DACOMP tended to be the largest among all competing methods, it is significantly lower than that of LOCOM. WRENCH is only applicable to one scenario (with a binary trait and no confounder) in which case it had inflated FDR and nevertheless low sensitivity.

Results for simulated data with differential experimental bias (and a binary trait and no confounder) are shown in Figure 5. These simulations showed that while LOCOM, ANCOM, and DACOMP were unaffected by differential bias, ANCOM-BC, ALDEx2, and WRENCH were sensitive to differential bias, and yielded significantly inflated FDR in the presence of such bias.

**Figure 5:**
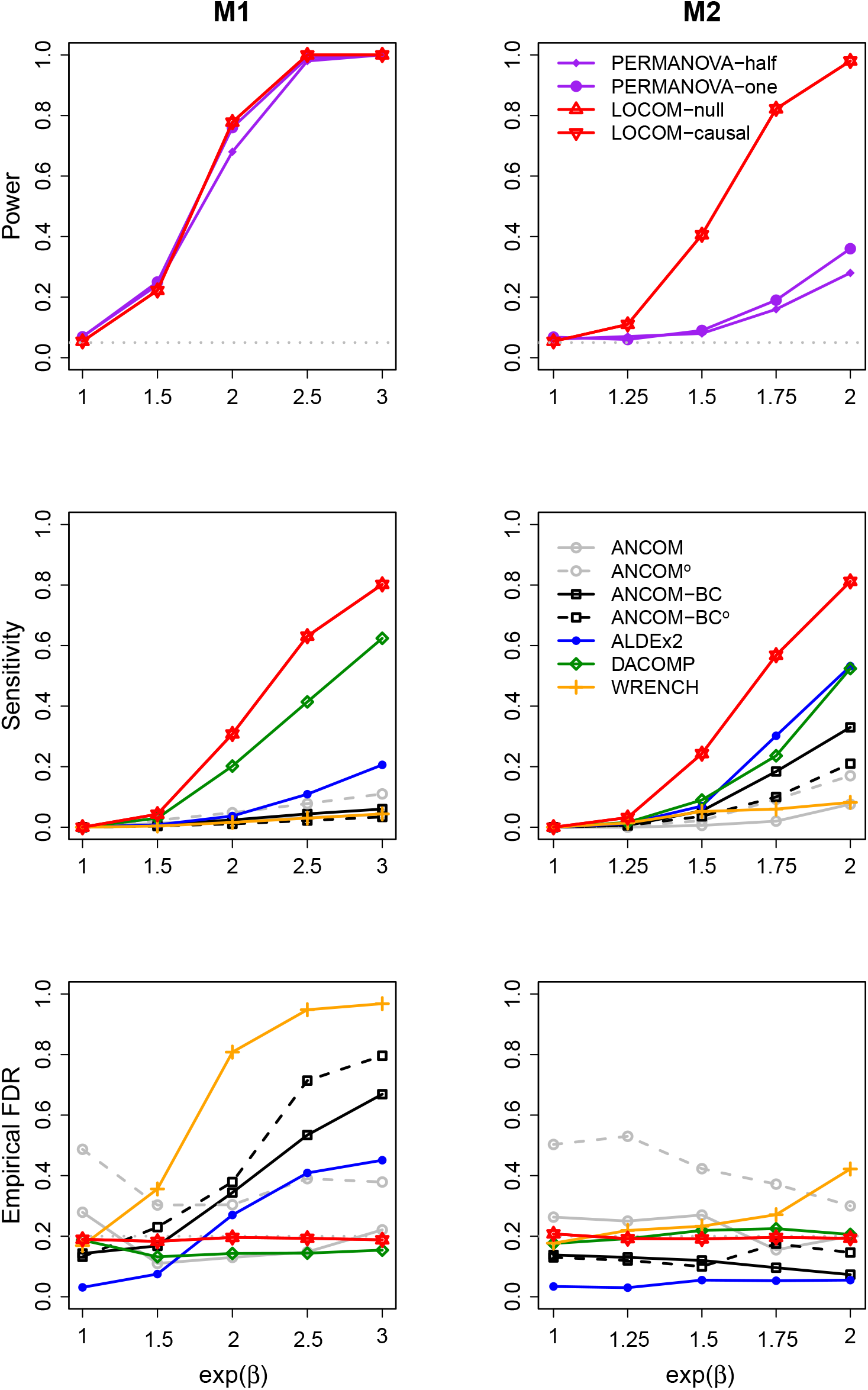
Simulation results for data (*n* = 100) with differential experimental bias in the binary-trait setting (no confounder).

Results for simulated data with heterogeneous *β*_*j*,1_ values are displayed in Figure S9. The patterns we observed with heterogeneous *β*_*j*,1_ values were similar to those seen in the analogous simulations with homogeneous *β*_*j*,1_ values (Figure 3).

### URT microbiome data

The data for our first example were generated as part of a study to examine the effect of cigarette smoking on the orpharyngeal and nospharyngeal microbiome [35]. We focused on the left orpharyngeal microbiome in this analysis. The 16S sequence data were summarized into a taxa count table consisting of data from 60 samples and 856 taxa. The trait of interest was a binary variable for smoking status, which classified the samples into 28 smokers and 32 nonsmokers. Other covariates include gender and antibiotic use within the last 3 months. There was an imbalance in the proportion of males by smoking status (75% in smokers, 56% in non-smokers), indicating a potential confounding effect of gender. Since there were only three samples who used antibiotics within the last 3 months, we excluded these samples from our analysis and adjusted for gender only. We adopted the same filter (20% presence) as in the simulation studies, which resulted in 111 taxa for downstream analysis. We applied LOCOM with the most abundant taxon (having mean relative abundance 10.5% before filtering and 11.4% after filtering) as the reference. Given the need to adjust for gender, we only applied ANCOM, ANCOM-BC, and ALDEx2 as a comparison. The nominal FDR was set at 10%.

As shown in the upper panel of Table 2, the global *p*-value of LOCOM is 0.0045, which indicates a significant difference in the overall microbiome profile between smokers and non-smokers after adjusting for gender. At the taxon level, LOCOM, ALDEx2, ANCOM, and ANCOM-BC detected 6, 0, 2, and 2 taxa, respectively; Figure 6 displays a Venn diagram of these sets of taxa; Table S2 lists information on the 6 taxa detected by LOCOM. Figure 7 shows the distributions of relative abundance across four covariate groups cross-classified by smoking status and gender, for taxa detected by LOCOM, ANCOM, and ANCOM-BC, as well as for two null taxa. One null taxon is the taxon with the median 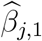 value. The other is the average of a group of null taxa for improved stability. The two null taxa both had lower relative abundance in smokers than in non-smokers, among either females or males. The six taxa detected by LOCOM all had the opposite trend (i.e., higher relative abundance in smokers than in non-smokers), indicating that these taxa are likely to be real signals (i.e., overgrew in smokers). The taxon detected by ANCOM only also had the opposite trend to the null taxa, but it was not detected by LOCOM because the adjusted *p*-value (0.13) by LOCOM did not meet the nominal FDR. The taxon detected by ANCOM-BC only had a similar trend as the null taxa, suggesting that this taxon may actually be a null taxon; indeed, the adjusted *p*-value by LOCOM is 0.689. Note that the difference in relative abundance distributions between smokers and non-smokers at null taxa may be considered as the counterbalancing change that the null taxa underwent in response to the changes at the causal taxa.

**Table 2:** Results in analysis of the two real datasets

| Trait | Global $p$-value | Number of detected taxa | | | |
| --- | --- | --- | --- | --- | --- |
|  | LOCOM | LOCOM | ALDEx2 | ANCOM | ANCOM-BC |
| URT microbiome data |  |  |  |  |  |
| Smoking | 0.0045 | 6 | 0 | 2 | 2 |
| PPI microbiome data |  |  |  |  |  |
| gPC1 | 0.70 | 0 | 0 | NA | NA |
| gPC2 | 0.020 | 2 | 0 | NA | NA |
| gPC3 | 0.018 | 2 | 0 | NA | NA |
| gPC4 | 0.16 | 0 | 0 | NA | NA |
| gPC5 | 0.0070 | 32 | 0 | NA | NA |
| gPC6 | 0.59 | 0 | 0 | NA | NA |
| gPC7 | 0.11 | 0 | 0 | NA | NA |
| gPC8 | 0.21 | 0 | 0 | NA | NA |
| gPC9 | 0.11 | 0 | 0 | NA | NA |

Note: ANCOM and ANCOM-BC are not applicable for testing continuous traits.

**Figure 6:**
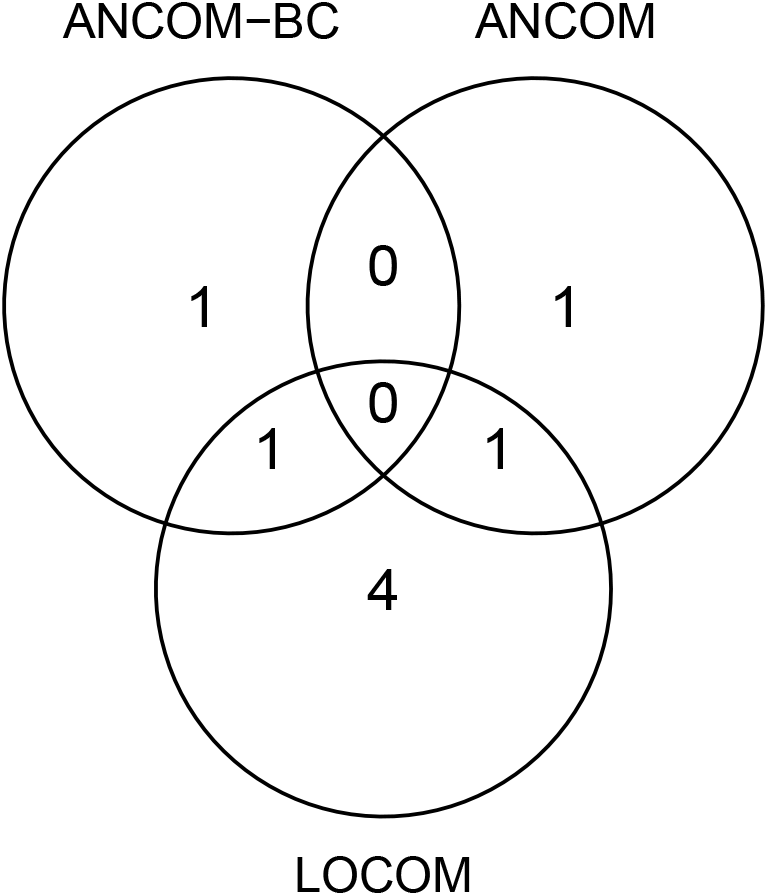
Taxa detected to be differentially abundant in the URT data.

**Figure 7:**
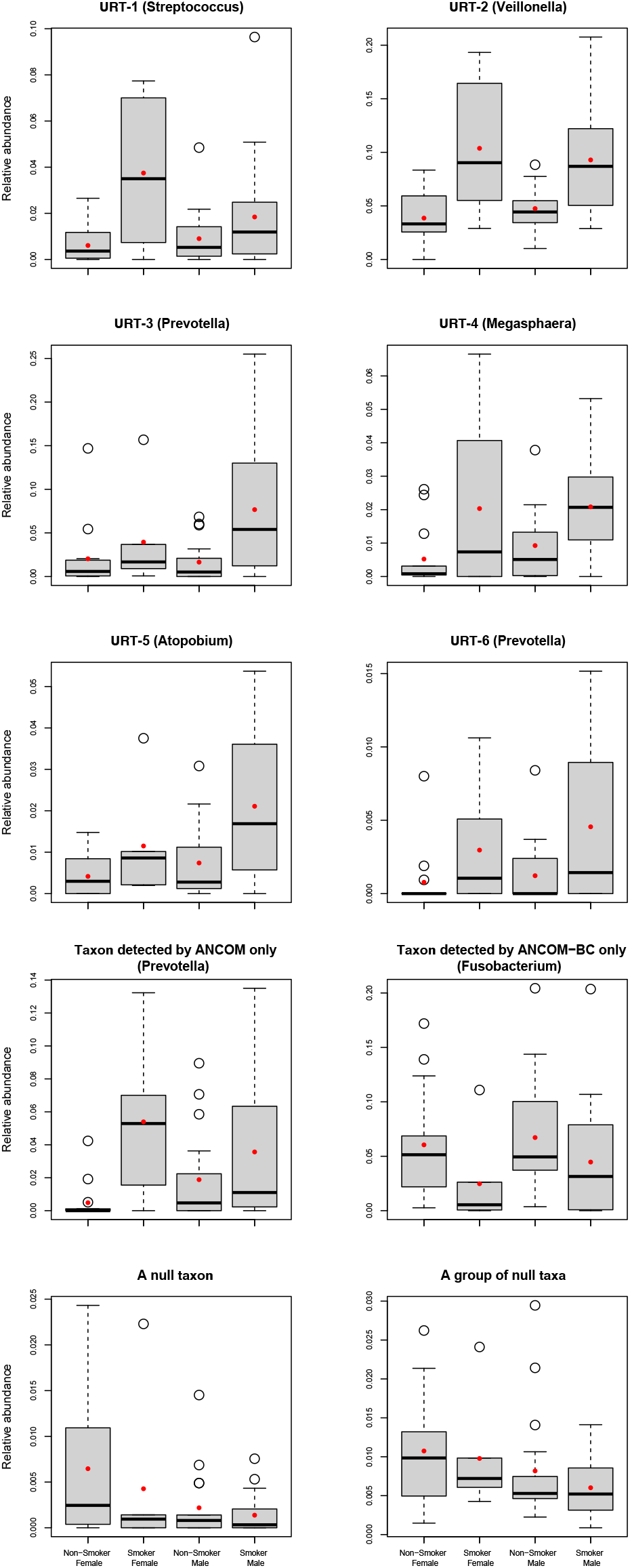
Distributions of relative abundances for taxa in the URT data. The six taxa in rows 1-3 were detected by LOCOM; among these, URT-2 was also detected by ANCOM-BC and URT-3 was also detected by ANCOM. In the last row, “A null taxon” corresponds to the taxon (*Shigella*) with the median 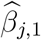 value. “A group of null taxa” include the taxon with the median 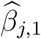 value and 20 taxa with 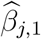 values closest to (10 less than and 10 greater than) the median; their relative abundances were averaged.

The original analysis of this dataset [35] reported that *Megasphaera* and *Veillonella spp*. were most enriched in the left oropharynx of smokers compared to non-smokers. Later, a large study of oral microbiome (from oral wash samples) in 1204 American adults [36] reported enrichment of *Atopobium, Streptococcus*, and *Veillonella* in smokers compared to non-smokers. More recently, a shotgun metagenomic sequencing study of salivary microbiome in Hungary population [37] reported enrichment of *Prevotella* and *Megasphaera* in smokers compared to non-smokers. Thus, all six taxa detected by LOCOM have been implicated in the literature, even if we only consider the latter two independent studies. These taxa were largely missed by ANCOM and ANCOM-BC.

### PPI microbiome data

The data for our second example were generated in a study of the association between the mucosal microbiome in the prepouch-ileum (PPI) and host gene expression among patients with IBD [38]. The PPI microbiome data from 196 IBD patients were summarized in a taxa count table with 7,000 taxa classified at the genus level. The gene expression data at 33,297 host transcripts, as well as clinical metadata such as antibiotic use (yes/no), inflammation score (0–9), and disease type (familial adenomatous polyposis/FAP and non-FAP) were also available. The data also included nine gene principal components (gPCs) that together explained 50% of the total variance in host gene expression. Here, we included all nine gPCs as multiple traits of interest into one model while adjusting for the three potentially confounding covariates. We filtered out taxa based on our previous filtering criterion, which resulted in 507 taxa to be included in the analysis. We applied LOCOM with the most abundant (8.2%) taxon as the reference. Given the continuous traits of interest and the three covariates, we only considered ALDEx2 for comparison. The nominal FDR was set at 10%.

The results of PPI data analysis are presented in the lower panel of Table 2. LOCOM discovered that gPC2, gPC3, and gPC5 had significant associations with the overall microbial profiles at the *α* = 0.05 level. LOCOM detected 2, 2, and 32 taxa as associated with gPC2, gPC3, and gPC5, respectively, at the 10% FDR level, and did not detect any taxa for the gPCs that were not found to be associated with the microbiome by the global test. Among the 32 taxa associated with gPC5, 15 belong to the genus *Escherichia* (Table S2), which appeared frequently in the literature of IBD according to a highly-cited review article [39]. ALDEx2 failed to detect any taxa.

## Discussion

We have presented LOCOM, a novel compositional approach for testing differential abundance in the microbiome data, at both the taxon level and the global level. The global statistic is an aggregate of *p*-values from tests of individual taxa, so results from the taxon-level and global tests are coherent. LOCOM allows both binary and continuous traits of interest, can test multiple traits simultaneously, and can adjust for confounding covariates. In our simulations, the taxa detected by LOCOM always preserved FDR while those identified by the competing methods did not, even though LOCOM had clearly superior sensitivity. In addition, LOCOM also provided a global test that always controled the type I error and had good power compared to PERMANOVA. In analysis of the URT microbiome data, we demonstrated that the taxa detected by LOCOM were likely to be real signals while those detected by ANCOM and/or ANCOM-BC but not LOCOM may be false positives. In analysis of the PPI microbiome data, since global and taxon-specific tests were coherent, LOCOM identified significant taxa only for gene principal components that were globally significant.

It is possible to generalize LOCOM to test a categorical trait with more than two levels. Ordered categories could be handled in the framework presented here by assigning an appropriate score to each category and then treating this score as a continuous variable. For a categorical trait with *K* unordered categories, we would presumably need to estimate *K* − 1 effect sizes to fully describe the variable; we could then compare some summary (e.g., max or mean) of these effect sizes to the equivalent value in the null permutations. Although this better analysis would require some software development and simulation testing, a simpler proposal could provide results within the existing framework, by calculating separate (marginal) *p*-values for each of the *K* − 1 components and then combining these *p*-values into a single test statistic, e.g., by using the harmonic mean statistic we used to form our global test. Choosing these *K* − 1 components to be orthogonal may be helpful here. We hope to modify LOCOM to incorporate multi-category variables in future work.

Our filtering criterion to exclude taxa with fewer than 20% presence in the sample worked well for the extensive simulation studies we conducted. In fact, a compositional analysis performs best when non-null taxa are relatively common throughout all samples. Analyses that look for the effect of rare taxa should probably be focussed on a presence-absence analysis [40, 41], or on a method based directly on relative abundances.

The compositional null hypothesis considered here is also appropriate in other experimental settings, such as studies of gene expression. This hypothesis corresponds to the scenario that a small number of microbes have “bloomed” while the absolute counts of the others have not changed; this is the reason we made the assumption that more than half of the taxa are null taxa, which is commonly made in other compositional methods. In the gene expression experiment, we often see only a few genes that are differentially expressed; the majority of genes have the same expression in cases and controls. However, it is not completely clear that the compositional hypothesis is applicable to microbiome data because, unlike genes, microbes interact with each other: not only do they compete for resources, but they also change their environment in ways that favor some microbes and suppress others. For example, *Lactobacilli* generally make lactic acid, which changes the pH of the environment. This suppresses microbes that do not thrive in an acidic environment while encouraging growth of microbes that do. Because the microbiota are a community, it is not unreasonable to expect that potentially every taxon changes between cases and controls. The “community change” null hypothesis may also be reasonable because, when comparing the alpha diversity with causal taxa spiked in to a case group, the control group would have a lower alpha diversity (i.e., lower evenness); if this change in alpha diversity is meaningful, then the “community change” null hypothesis is appropriate. When the “community change” null hypothesis seems more reasonable than the compositional null hypothesis, then a method that applies directly to relative abundance data such as the LDM is more appropriate. Note that, unlike the compositional null, the “community change” null hypothesis will consider *all* taxon relative abundances to be potentially changed if extra counts of a small number of taxa are “spiked in”. However, the LDM when applied to relative abundance data is not invariant to experimental bias the way LOCOM is; in fact, hypotheses based on differences in relative abundances typically require tests based on unbiased data to be valid.

We showed both theoretically and with simulation studies that LOCOM is unaffected by experimental bias, even when bias factors are differentially distributed between causal and non-causal taxa. While some competing compositional methods (ANCOM and DACOMP) share this robustness, others (ANCOM-BC, ALDEx2 and WRENCH) do not. This may be related to the choice of centering; in general, the centered log ratio will not be robust when there are cells with zero counts, since this centering will depend on the set of taxa seen in each sample even if a pseudocount is used. Thus, the centering may not cancel out when comparing log ratios from different samples, leaving these comparisons affected by the particular bias factors that characterize the data being analyzed. Note that any compositional method should perform well when the bias is non-differential, since the centering will be the same on average in each sample.

We have implemented our method in the R package LOCOM, which is computationally efficient for data with small sample sizes but can take longer for larger sample sizes. For example, using a single-thread MacBook Pro laptop (1.4 GHz Quad-Core Intel Core i5, 8GB memory), it took 80s and 120s to analyze a simulated dataset with 100 and 200 samples, respectively, 14.4s to analyze the URT data, but 2 days to analyze the PPI data. In considering this last timing, it should be noted that the analysis considered 9 traits simultaneously in the presence of 3 confounding covariates, and as such is more complex than the typical microbiome analysis. In addition, LOCOM can be easily parallelized by splitting the permutation replicates into different machines and then counting the total number of rejection, and further by splitting the data into subsets with sets of taxa that only share the reference taxon and then combining the values of *β*_*j*,1_ from each dataset (care should be taken to use the same seed for each analysis so that the same set of permutations is used).

