## Supplemental Figures and Tables for "LOCOM: A logistic regression model for testing differential abundance in compositional microbiome data with false discovery rate control"

**Table S1:** Type I error for testing the global hypothesis at nominal level 0.05, without adjustment of the confounder

| Trait | Confounder | Causal mechanism | LOCOM -null | LOCOM -causal |
|---|---|---|---|---|
| Binary | Binary | M1 | 0.132 | 0.132 |
| | | M2 | 0.154 | 0.154 |
| Binary | Continuous | M1 | 0.481 | 0.481 |
| | | M2 | 0.525 | 0.525 |
| Continuous | Binary | M1 | 0.161 | 0.145 |
| | | M2 | 0.168 | 0.177 |
| Continuous | Continuous | M1 | 0.112 | 0.104 |
| | | M2 | 0.123 | 0.139 |

Note: Sample size $n = 100$.

**Table S2:** Taxa (in ascending order of the raw $p$-value) detected by LOCOM in analysis of the two real datasets

| Taxon ID | Taxon Assignment | Mean relative abundance | Raw $p$-value | Adjusted $p$-value |
|---|---|---|---|---|
| Taxa associated with smoking in URT microbiome data | | | | |
| URT-1 | *g_Streptococcus* | 0.015 | 0.00036 | 0.028 |
| URT-2 | *g_Veillonella* | 0.067 | 0.00055 | 0.028 |
| URT-3 | *g_Prevotella* | 0.041 | 0.0015 | 0.033 |
| URT-4 | *g_Megasphaera* | 0.013 | 0.0015 | 0.033 |
| URT-5 | *g_Atopobium* | 0.012 | 0.0016 | 0.033 |
| URT-6 | *g_Prevotella* | 0.0024 | 0.0029 | 0.049 |
| Taxa associated with gPC5 in PPI microbiome data | | | | |
| PPI-1 | *f_Flavobacteriaceae* | 0.00098 | 0.0020 | 0.025 |
| PPI-2 | *g_Escherichia* | 0.015 | 0.0020 | 0.025 |
| PPI-3 | *g_Escherichia* | 0.00032 | 0.0020 | 0.025 |
| PPI-4 | *f_Comamonadaceae* | 0.0013 | 0.0040 | 0.030 |
| PPI-5 | *g_Coprococcus* | 0.00051 | 0.0040 | 0.030 |
| PPI-6 | *g_Escherichia* | 0.093 | 0.0060 | 0.032 |
| PPI-7 | *g_Escherichia* | 0.0021 | 0.0060 | 0.032 |
| PPI-8 | *f_Enterobacteriaceae* | 0.00037 | 0.0014 | 0.066 |
| PPI-9 | *g_Blautia* | 0.00015 | 0.0018 | 0.070 |
| PPI-10 | *g_Acinetobacter* | 0.0014 | 0.0022 | 0.070 |
| PPI-11 | *g_Escherichia* | 0.0011 | 0.0022 | 0.070 |
| PPI-12 | *g_Streptococcus* | 0.00023 | 0.0024 | 0.070 |
| PPI-13 | *g_Escherichia* | 0.0032 | 0.0026 | 0.070 |
| PPI-14 | *g_Escherichia* | 0.000058 | 0.0026 | 0.070 |
| PPI-15 | *g_Escherichia* | 0.00028 | 0.0028 | 0.070 |
| PPI-16 | *g_Escherichia* | 0.0053 | 0.0032 | 0.075 |
| PPI-17 | *g_Escherichia* | 0.000071 | 0.0034 | 0.075 |
| PPI-18 | *Unclassified* | 0.00091 | 0.0044 | 0.085 |
| PPI-19 | *g_Streptococcus* | 0.00010 | 0.0046 | 0.085 |
| PPI-20 | *g_Bifidobacterium* | 0.011 | 0.0052 | 0.085 |
| PPI-21 | *f_Clostridiaceae* | 0.0015 | 0.0054 | 0.085 |
| PPI-22 | *g_Escherichia* | 0.00016 | 0.0056 | 0.085 |
| PPI-23 | *g_Escherichia* | 0.000080 | 0.0058 | 0.085 |
| PPI-24 | *g_Escherichia* | 0.00011 | 0.0062 | 0.085 |
| PPI-25 | *g_Bifidobacterium* | 0.0015 | 0.0064 | 0.085 |
| PPI-26 | *f_Lachnospiraceae* | 0.000089 | 0.0064 | 0.085 |
| PPI-27 | *g_Escherichia* | 0.000049 | 0.0064 | 0.085 |
| PPI-28 | *g_Clostridium* | 0.00031 | 0.0066 | 0.085 |
| PPI-29 | *g_Blautia* | 0.0040 | 0.0066 | 0.085 |
| PPI-30 | *g_Escherichia* | 0.00080 | 0.0068 | 0.085 |
| PPI-31 | *f_Enterobacteriaceae* | 0.000040 | 0.0072 | 0.088 |
| PPI-32 | *f_Enterococcaceae* | 0.00076 | 0.0080 | 0.094 |
| Taxa associated with gPC2 in PPT microbiome data | | | | |
| PPI-19 | *g_Streptococcus* | 0.00010 | 0.00020 | 0.047 |
| PPI-33 | *g_Streptococcus* | 0.00023 | 0.00020 | 0.047 |
| Taxa associated with gPC3 in PPT microbiome data | | | | |
| PPI-19 | *g_Streptococcus* | 0.00010 | 0.00020 | 0.041 |
| PPI-33 | *g_Streptococcus* | 0.00023 | 0.00020 | 0.041 |

Note: The mean relative abundance was calculated based on the data after filtering out taxa having fewer than 20% presence in the sample. "*g_*" or "*f_*" indicates that the taxon is assigned a genus or a family. The nominal FDR is 10%.

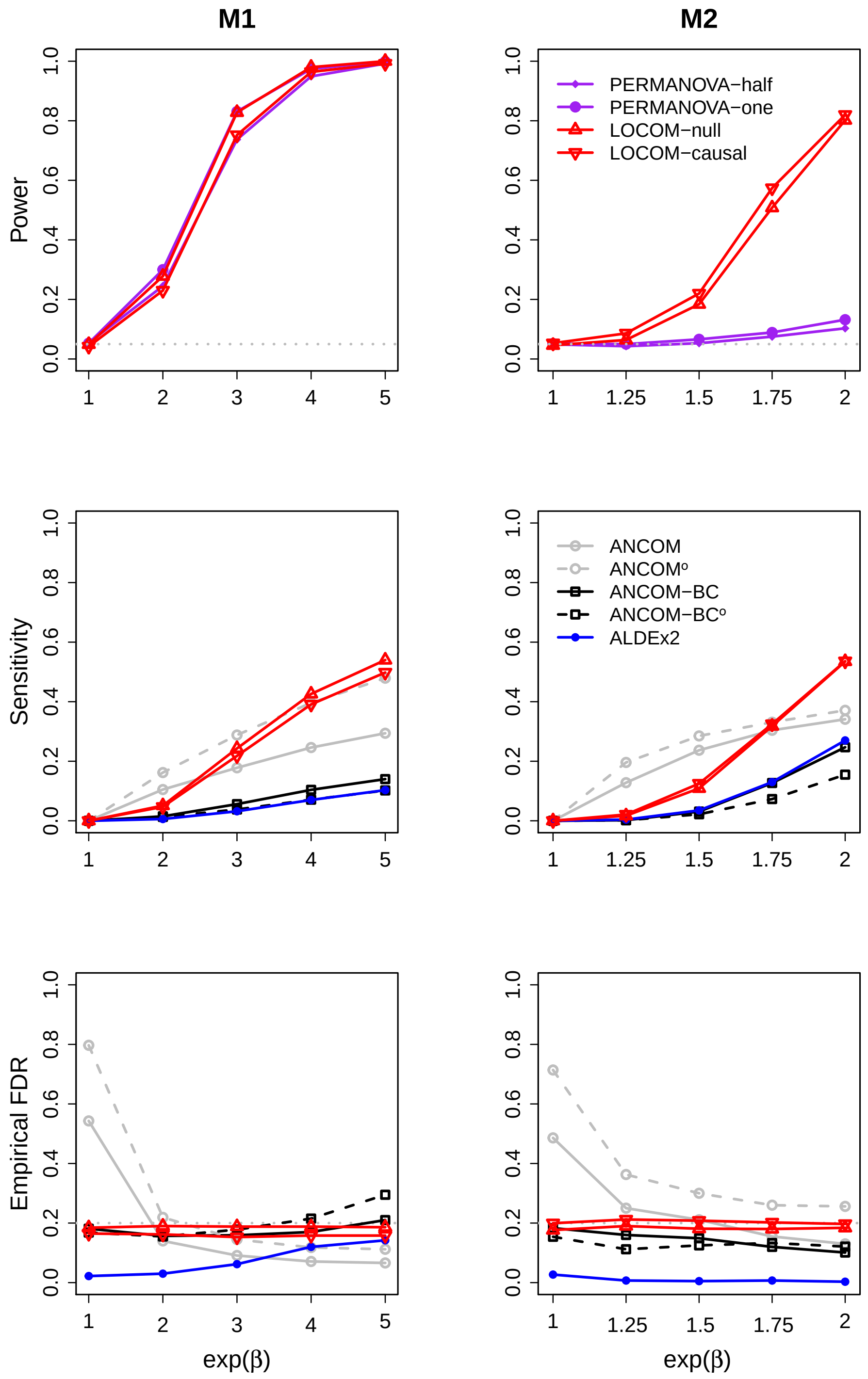


**Figure S1:** Simulation results for data ($n = 100$) with a binary trait and a continuous confounder.

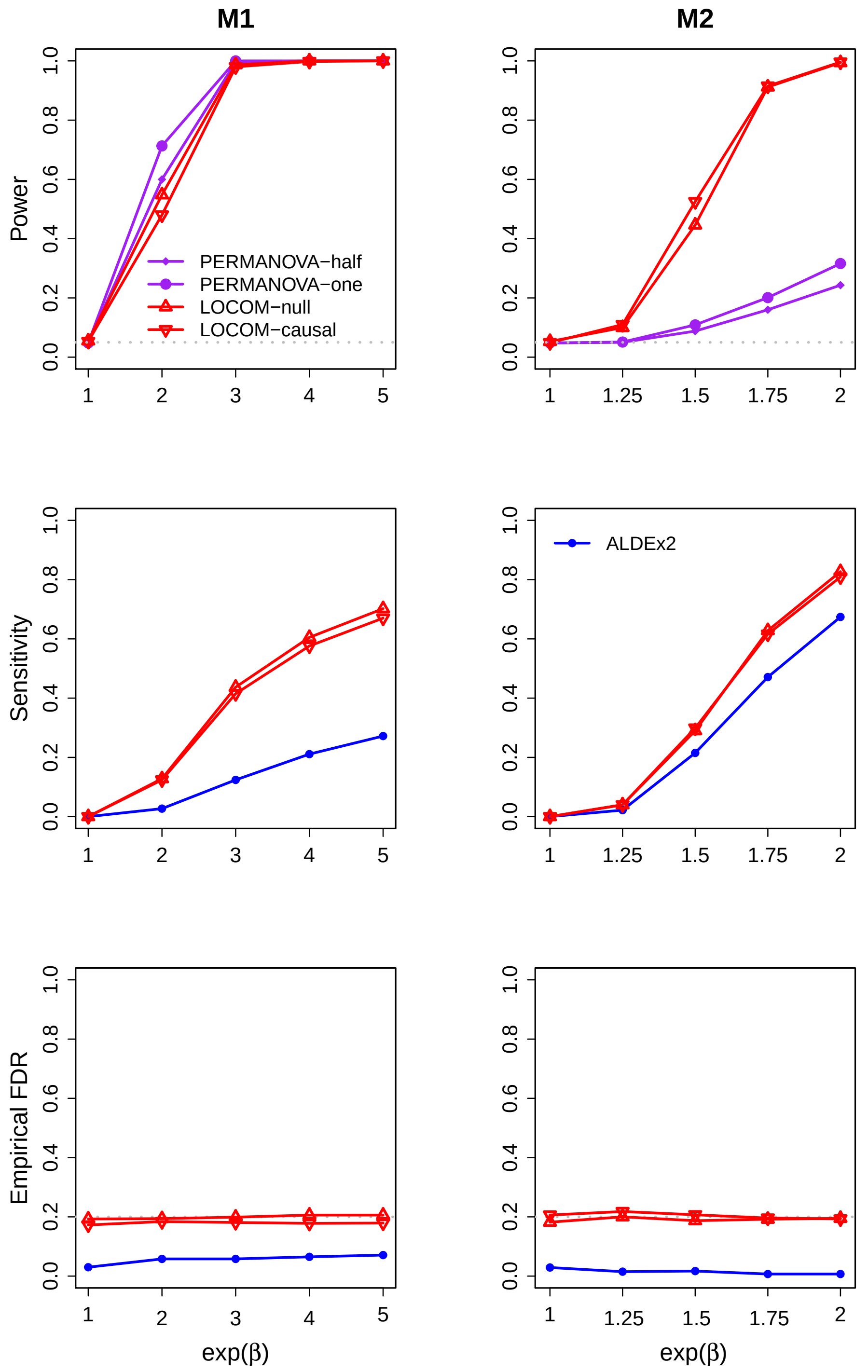


**Figure S2:** Simulation results for data ($n = 100$) with a continuous trait and a continuous confounder.

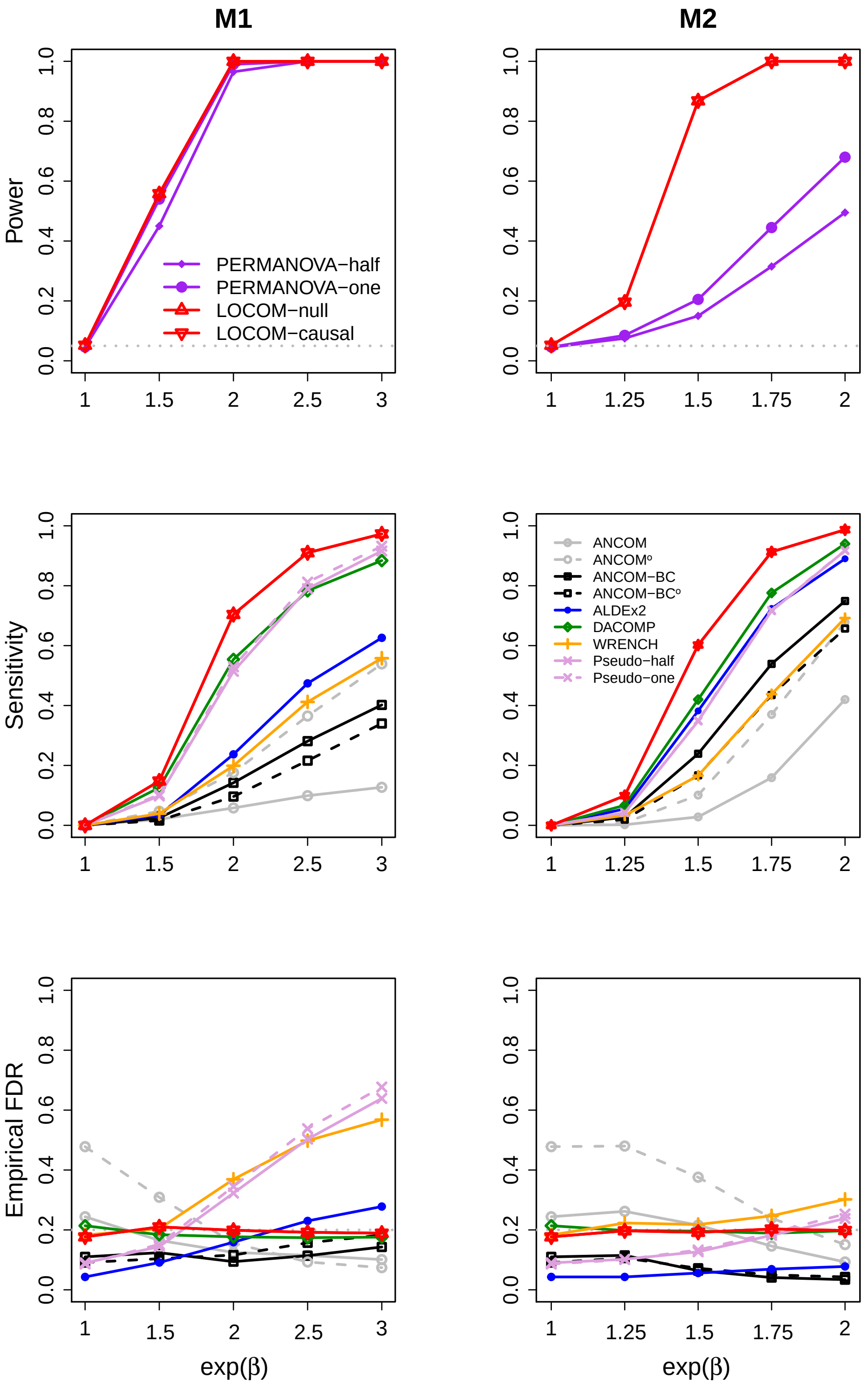


**Figure S3:** Simulation results for data ($n = 200$) with a binary trait (and no confounder).

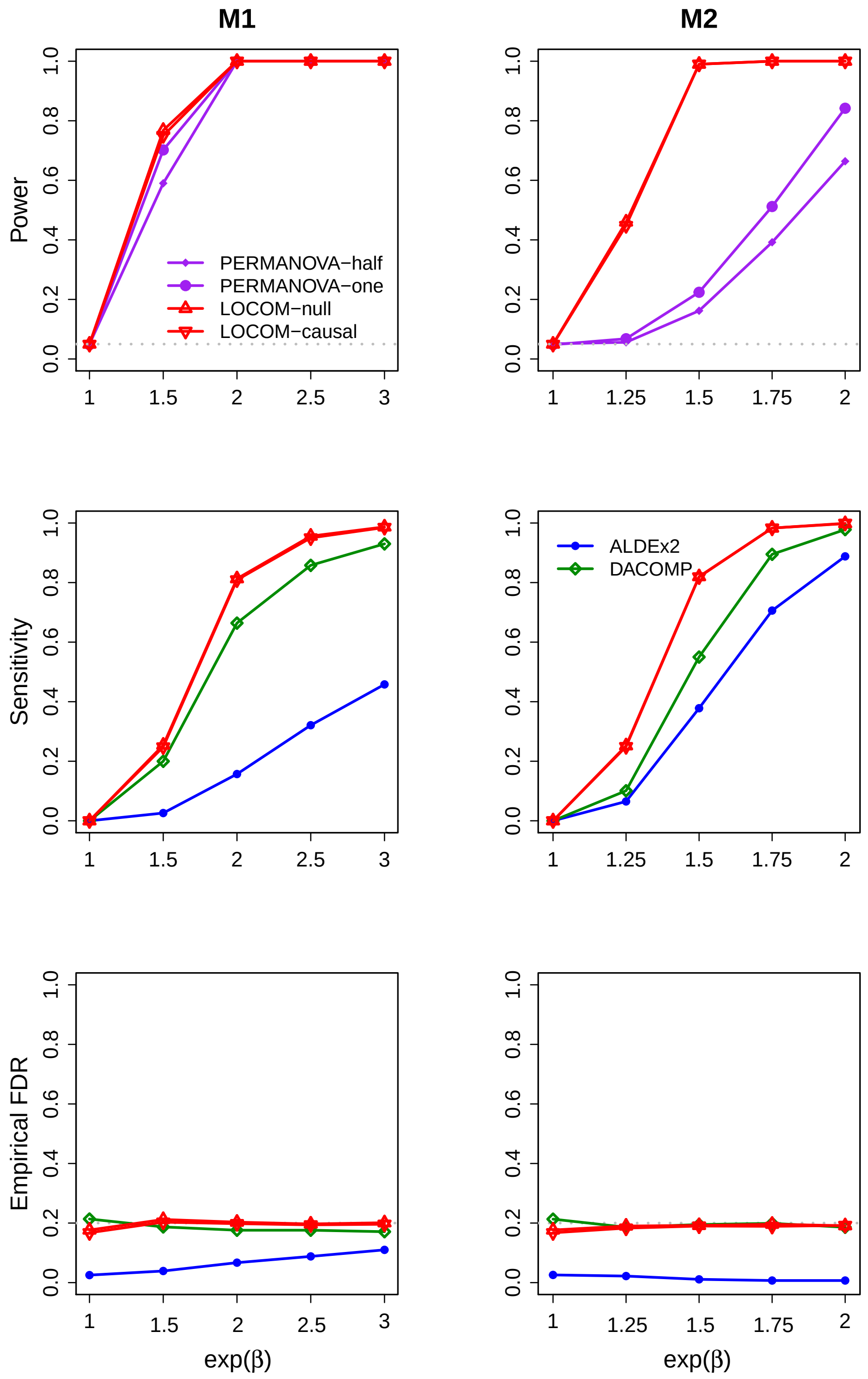


**Figure S4:** Simulation results for data ($n = 200$) with a continuous trait (and no confounder).

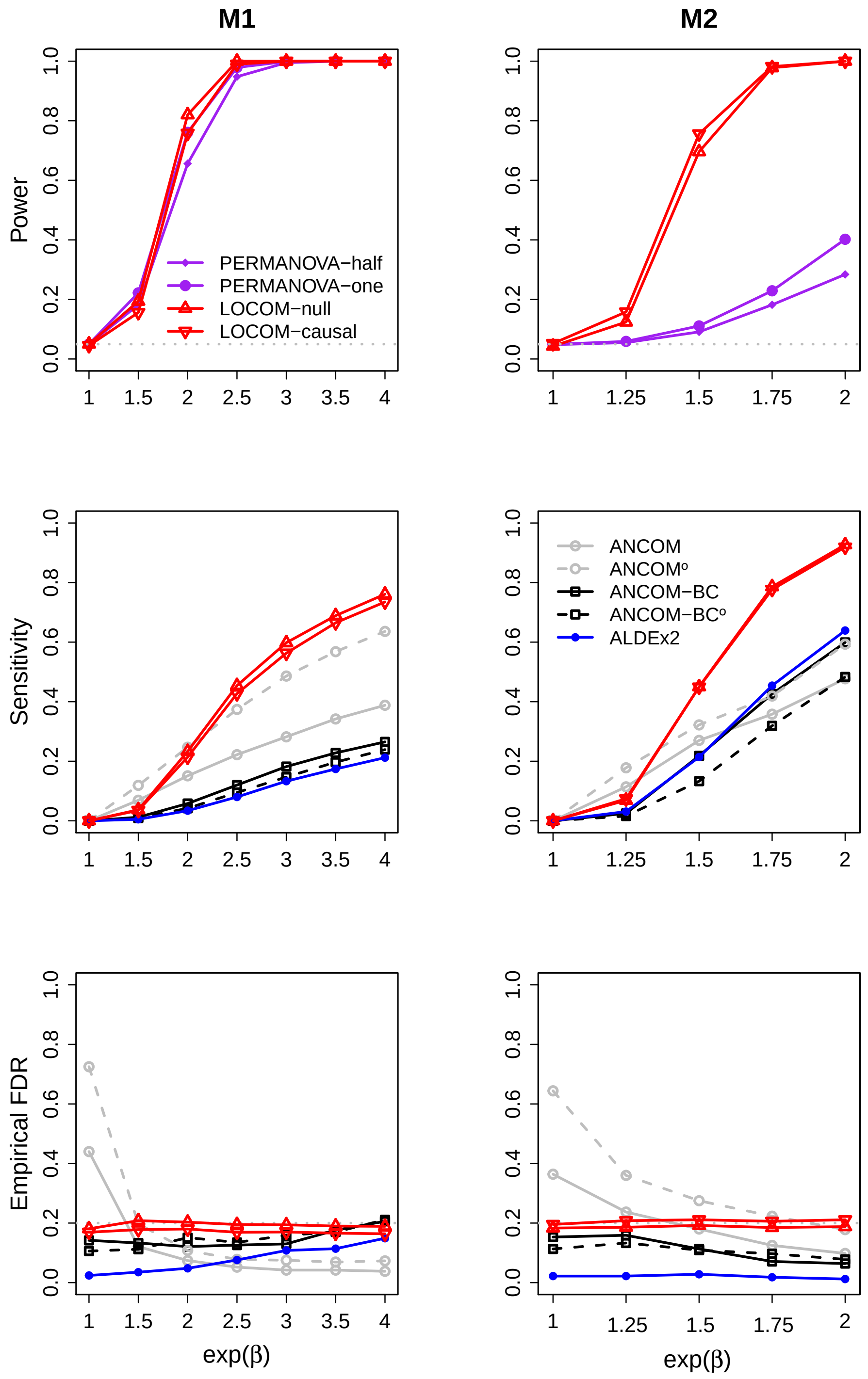


**Figure S5:** Simulation results for data ($n = 200$) with a binary trait and a binary confounder.

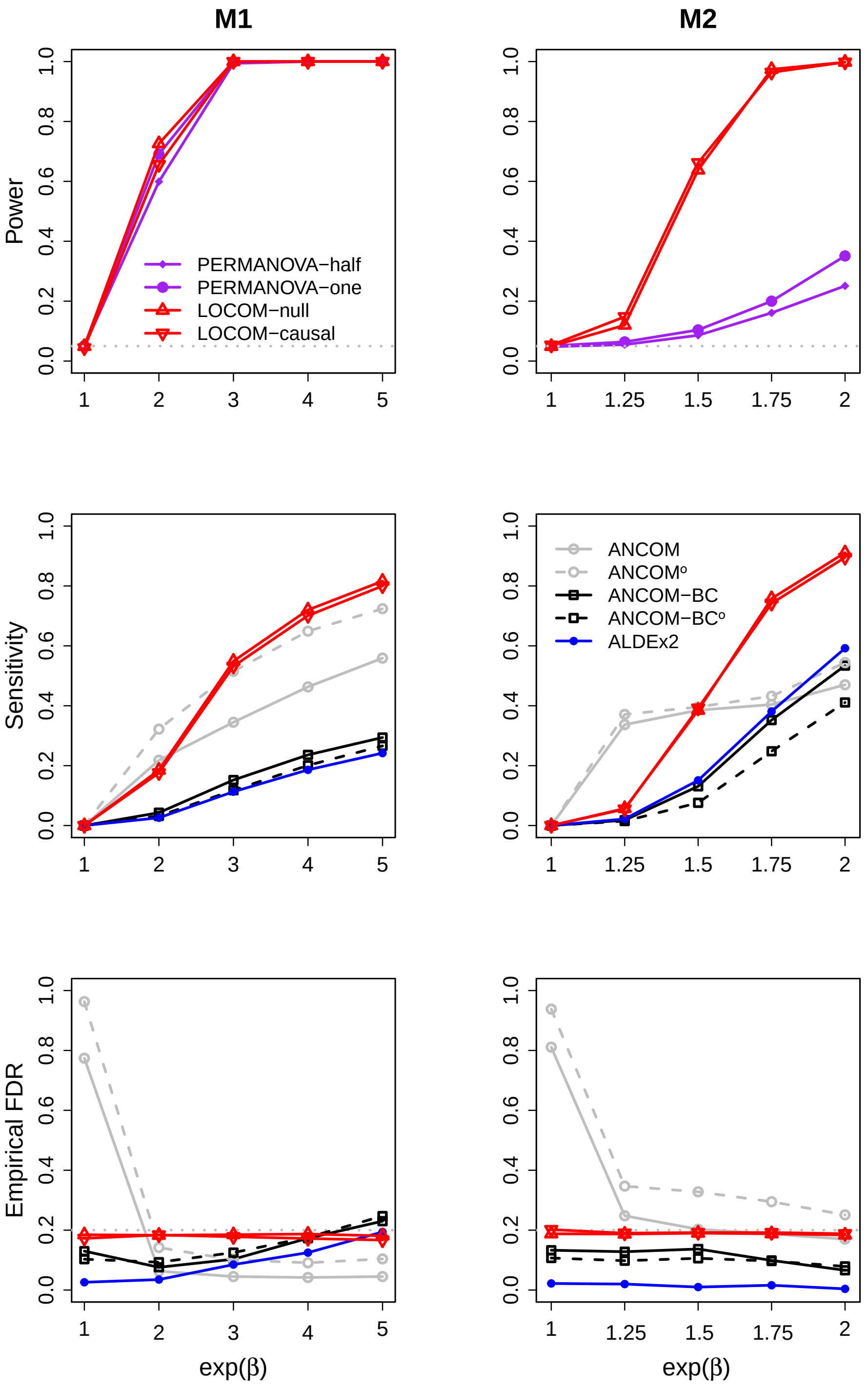


**Figure S6:** Simulation results for data ($n = 200$) with a binary trait and a continuous confounder.

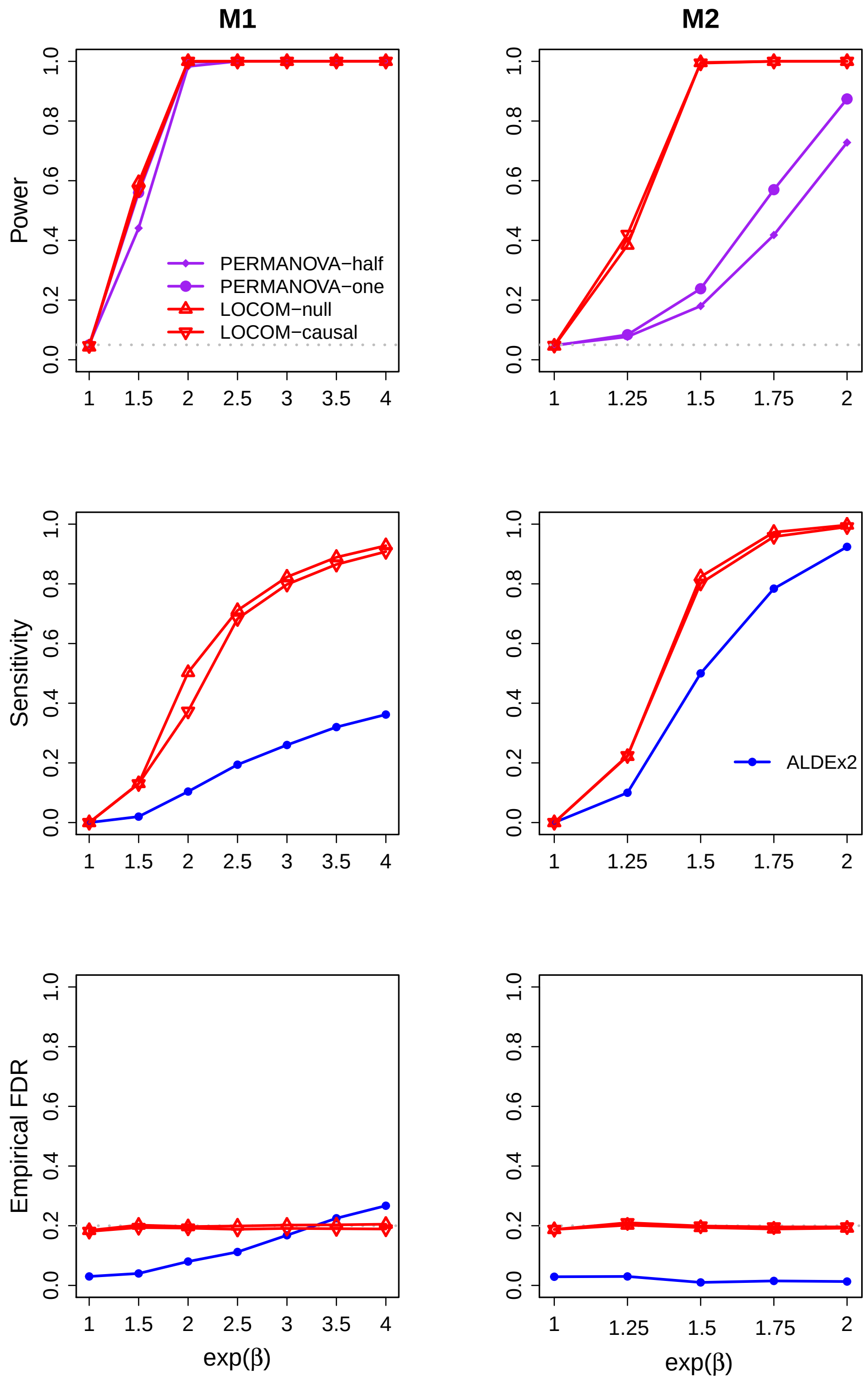


**Figure S7:** Simulation results for data ($n = 200$) with a continuous trait and a binary confounder.

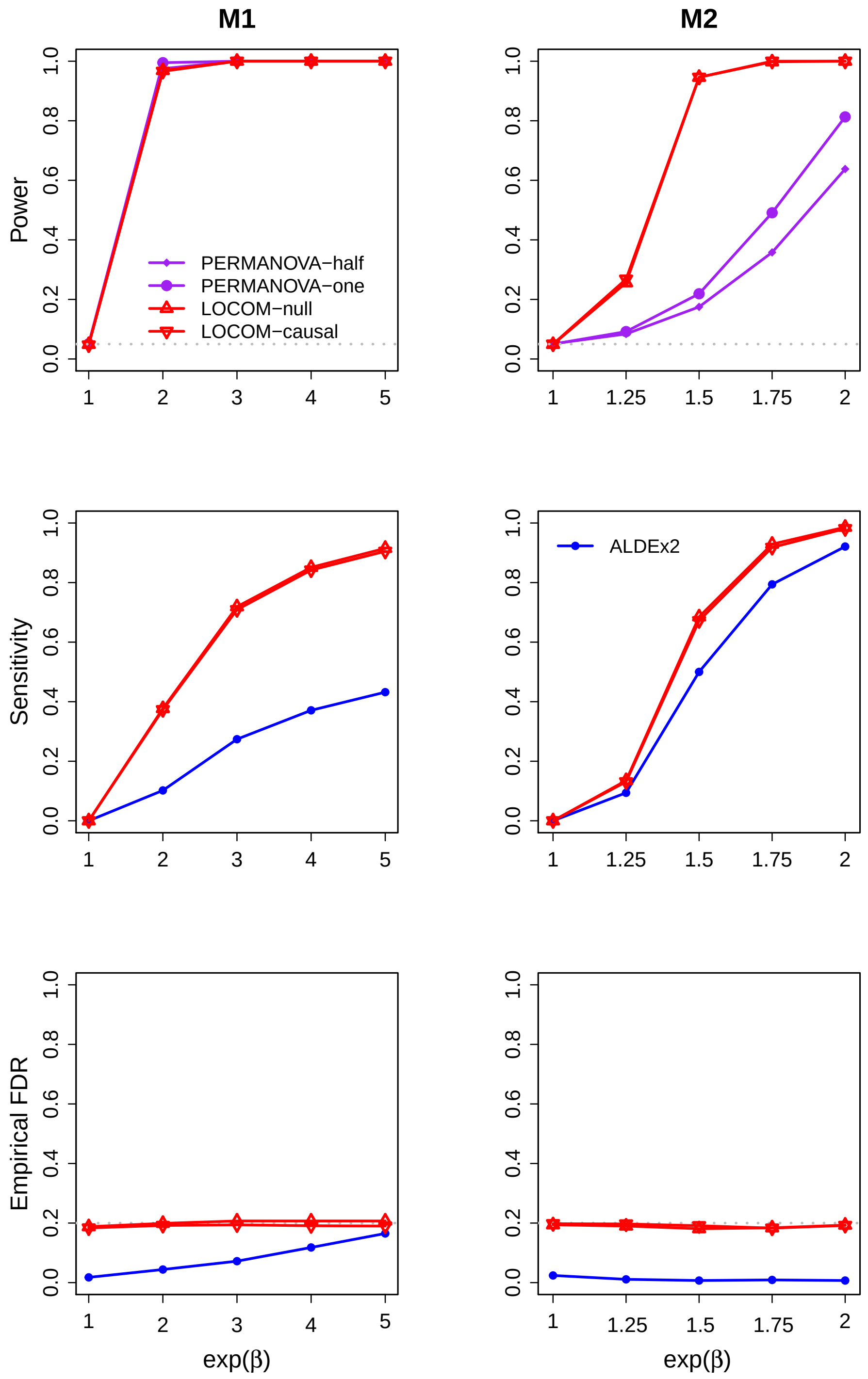


**Figure S8:** Simulation results for data ($n = 200$) with a continuous trait and a continuous confounder.

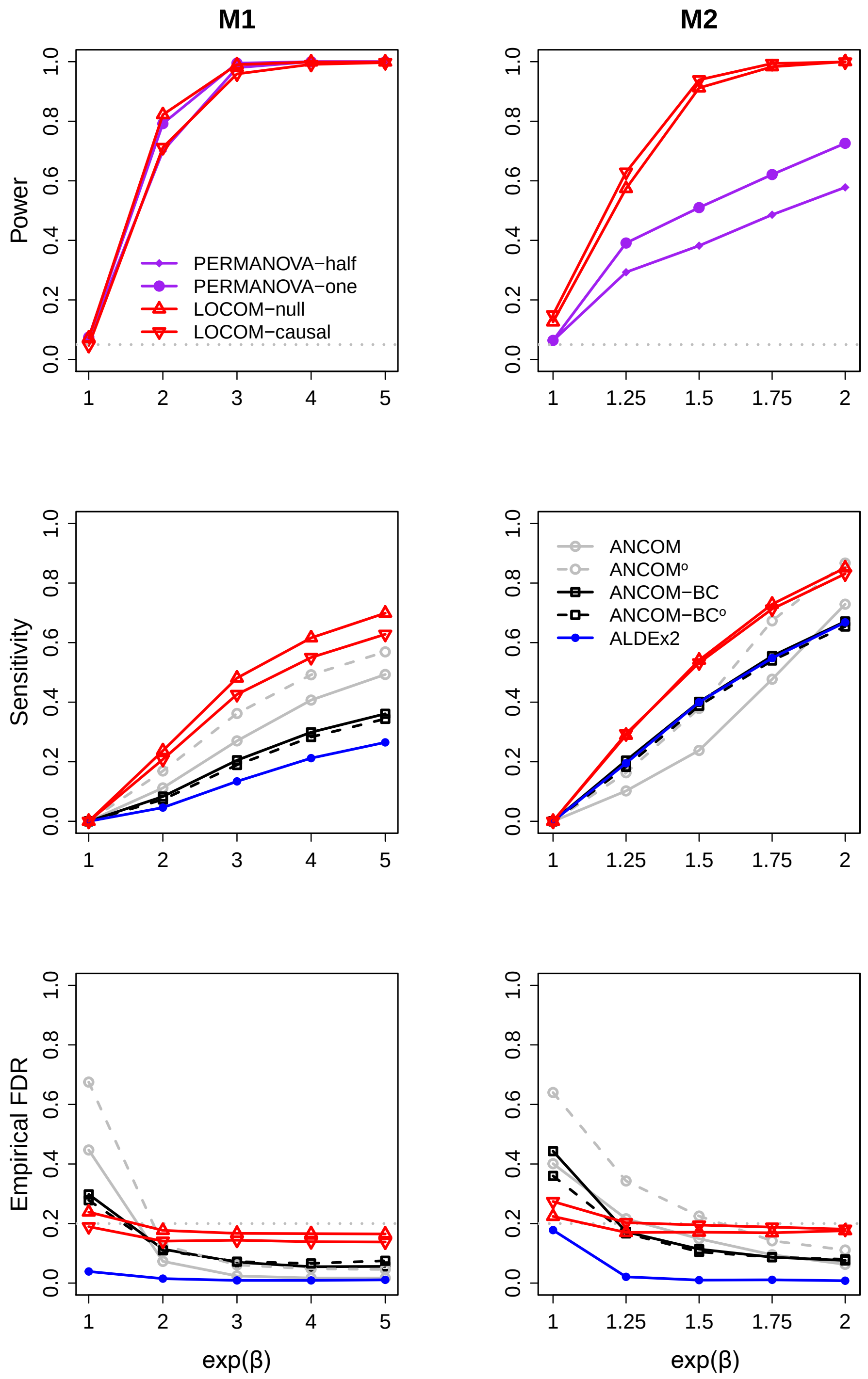


**Figure S9:** Simulation results for data ($n = 100$) with a binary trait and a binary confounder, when different values were sampled for different $\beta_{j,1}$, i.e., obtaining the fold change $\exp(\beta_{j,1})$ as an initial fold change scaled by the "scale factor". Specifically, the initial fold change was sampled from $U[0.5, 1.5]$, which implies different directions for initial fold changes of different taxa. If the initial fold change was positive (greater than 1), it was up-scaled (multiplied) by the scale factor to give the final fold change; if the initial fold change was negative (less than 1), it was down-scaled (divided) by the scale factor to give the final fold change.
